# Substrate-induced clustering activates Trim-Away of pathogens and proteins

**DOI:** 10.1101/2020.07.28.225359

**Authors:** Jingwei Zeng, Ana Filipa Santos, Aamir Mukadam, Mariana Osswald, Jakub Luptak, David Jacques, Claire Dickson, Nadine Renner, Chris Johnson, Marina Vaysburd, William A. McEwan, Eurico Morais-de-Sá, Dean Clift, Leo C. James

**Affiliations:** Medical Research Council, Laboratory of Molecular Biology, Cambridge CB2 0QH, United Kingdom; i3S - Instituto de Investigação e Inovação em Saúde, Universidade do Porto, Porto, Portugal; UK Dementia Research Institute, Department of Clinical Neurosciences, University of Cambridge, Cambridge, UK; EMBL Australia Node, Single Molecule Science, School of Medical Sciences, University of New South Wales, Sydney, Australia

**Keywords:** Trim-Away, TRIM21, Tripartite Motif, RING, E3 Ligase, ubiquitination, antibody, nanobody, molecular clustering, protein degradation, protein knockdown, optogenetics, PROTAC

## Abstract

Trim-Away is a powerful new technology that exploits off-the-shelf antibodies and the E3 RING ligase and cytosolic antibody receptor TRIM21 to carry out rapid protein depletion. How TRIM21 is catalytically-activated upon substrate engagement during either its normal immune function or when re-purposed for targeted protein degradation is unknown. Here we show that a mechanism of substrate-induced clustering triggers intermolecular dimerization of the RING domain to switch on the ubiquitination activity of TRIM21 and induce an antiviral response or drive Trim-Away. We harness this mechanism to expand the Trim-Away toolbox with highly-active TRIM21-nanobody chimeras that can also be controlled optogenetically. This work provides a mechanism for cellular activation of TRIM RING ligases and has important implications for targeted protein degradation technologies.

## INTRODUCTION

E3 ubiquitin ligases catalyse the final step in the ubiquitination reaction, which involves the transfer of ubiquitin from an E2 enzyme onto a substrate. There are more than 600 E3 ligases in the human genome, which regulate pathways involved in all aspects of eukaryotic biology. Typically, E3 ligases have evolved to ubiquitinate a dedicated substrate or set of substrates. Protein binding domains control which substrates a particular E3 ligase can interact with, while the E3 ligase active site modifies only those lysine residues that are correctly orientated to accept ubiquitin. Post-translational modifications such as phosphorylation and neddylation add further layers of regulation to control E3 ligase activation and ensure specificity of the ubiquitin transfer reactions (Zheng and Shabek, 2017). Sub-cellular localisation and tissue-specific expression provides the final level of regulation at the cell and organism level (Schapira et al., 2019; Yamada et al., 2013).

Despite this apparent exquisite specificity, E3 ligases are increasingly being re-purposed by researchers to target non-canonical substrates with ubiquitin. In this approach, intermediary molecules such as chemical compounds, peptides, antibodies or engineered binding domains are designed to artificially re-direct E3 ligases to proteins of interest, with the aim of causing their degradation through the ubiquitin-proteasome system. This approach has huge potential as a protein knockdown research tool as well as an entirely new therapeutic modality (Verma et al., 2020; Wu et al., 2020). However, precisely how E3 ligase activity is activated upon indirect recruitment to non-canonical substrates remains unclear.

Interestingly, evolution has already produced an E3 ubiquitin ligase whose function is to mediate degradation of diverse substrates via an intermediary molecule. TRIM21 is a member of the tripartite motif (TRIM) family of RING E3 ubiquitin ligases, but uniquely among TRIMs, and indeed all cytosolic proteins, TRIM21 possesses high-affinity antibody-binding activity (James et al., 2007). TRIM21 uses antibodies as intermediary molecules to target a wide-range of substrates for proteasomal degradation including viruses (Fan et al., 2016; Mallery et al., 2010; Vaysburd et al., 2013; Watkinson et al., 2015), bacteria (McEwan et al., 2013; Rakebrandt et al., 2014) and proteopathic agents such as Tau (McEwan et al., 2017). TRIM21 underpins a system of intracellular immunity where the diversity of the body’s antibody repertoire can be utilized in the cytosol to degrade invading pathogens (Foss et al., 2019; McEwan et al., 2011). This broad-spectrum targeting capability of TRIM21 has recently been exploited to develop Trim-Away, an easy to use and powerful method for acute and rapid degradation of endogenous cellular proteins. Trim-Away works by delivering antibodies to cells by electroporation or microinjection. Once in the cytosol, antibodies form a complex with both the target protein and TRIM21, leading to proteasomal degradation of the complex and a rapid protein depletion effect (Clift et al., 2017; Clift et al., 2018). Because Trim-Away utilizes standard off-the-shelf antibodies and no prior modification of proteins with tags or epitopes is required, this method has quickly been adopted to target a wide range of different proteins in various cell types and even model organisms (Castro-Dopico et al., 2019; Chen et al., 2019; Coyne et al., 2020; Fant et al., 2020; Israel et al., 2019; Nicolau et al., 2020; So et al., 2019).

Despite the important role of TRIM21 in the immune response and as a protein depletion technology, how TRIM21’s ubiquitin ligase activity is activated upon indirect substrate engagement via antibodies remains unknown. In this study we show that intermolecular clustering on the surface of an antibody-complexed substrate drives the dimerization of RING domains from neighbouring TRIM21 molecules and triggers K63-linked autoubiquitination. We exploit this mechanism to engineer highly active TRIM21-nanobody fusion proteins that combine protein targeting and catalytic activity into a single molecule that can be used to rapidly degrade cellular proteins. We further show that harnessing the clustering activation mechanism can be used to improve Trim-Away and allow precise optogenetic control of protein degradation.

## RESULTS

### TRIM21 assembles K63-linked ubiquitin chains on itself upon substrate engagement

TRIM21’s proteostatic functions are driven by its catalysis of ubiquitination, making control of this process a key point of cellular regulation. We therefore investigated whether substrate engagement is the trigger for TRIM21 enzymatic activity. We infected cells overexpressing His-tagged ubiquitin with adenovirus (AdV5) alone and in the presence of wild-type anti-Adv5 hexon antibody (9C12) or a mutant antibody that cannot bind TRIM21 [9C12(H433A)] (McEwan et al., 2012). TRIM21 did not undergo modification upon infection with Adv5, however in the presence of AdV5+9C12 TRIM21 underwent polyubiquitination. Polyubiquitination was lost when using the H433A antibody mutant, confirming that substrate engagement by TRIM21 is required (Figure 1A). Using cells over-expressing His-tagged TRIM21 but with endogenous ubiquitin, gave a similar, albeit weaker pattern of ubiquitination that was reversed with the deubiquitinase USP2 (Figure S1). To determine whether TRIM21 itself is activated, or if it merely becomes a substrate for another ubiquitin ligase, we introduced specific mutations into the TRIM21 RING domain that impair E2∼Ub binding (Figure 1B). These mutations, E12A/R/Y or E13A/R, significantly reduced polyubiquitination, with E12R and E13A showing the strongest effect (Figure 1A and consistent with their published impact in Trim-Away and Adv5 neutralization (Kiss et al., 2019). This indicates that TRIM21 is catalytically activated upon substrate binding. TRIM21 requires the K63-chain-specific E2 Ube2N for activity and to build anchored K63-chains *in vitro* on its own N-terminus after becoming primed with monoubiquitin (Fletcher et al., 2015). To address which ubiquitin chain types TRIM21 modifies itself with upon substrate engagement in cells, we expressed different ubiquitin mutants. Expression of K48R, which prevents K48-chain formation, did not impact TRIM21 polyubiquitination. Consistent with this, no polyubiquitination was seen in cells expressing a ubiquitin mutant that forms K48-linked chains only. In contrast, K63R substantially reduced TRIM21 polyubiquitination whilst cells expressing only K63 maintained chain formation (Figure 1D). This suggests that TRIM21 builds K63 chains on itself upon substrate engagement.

**Figure 1.**
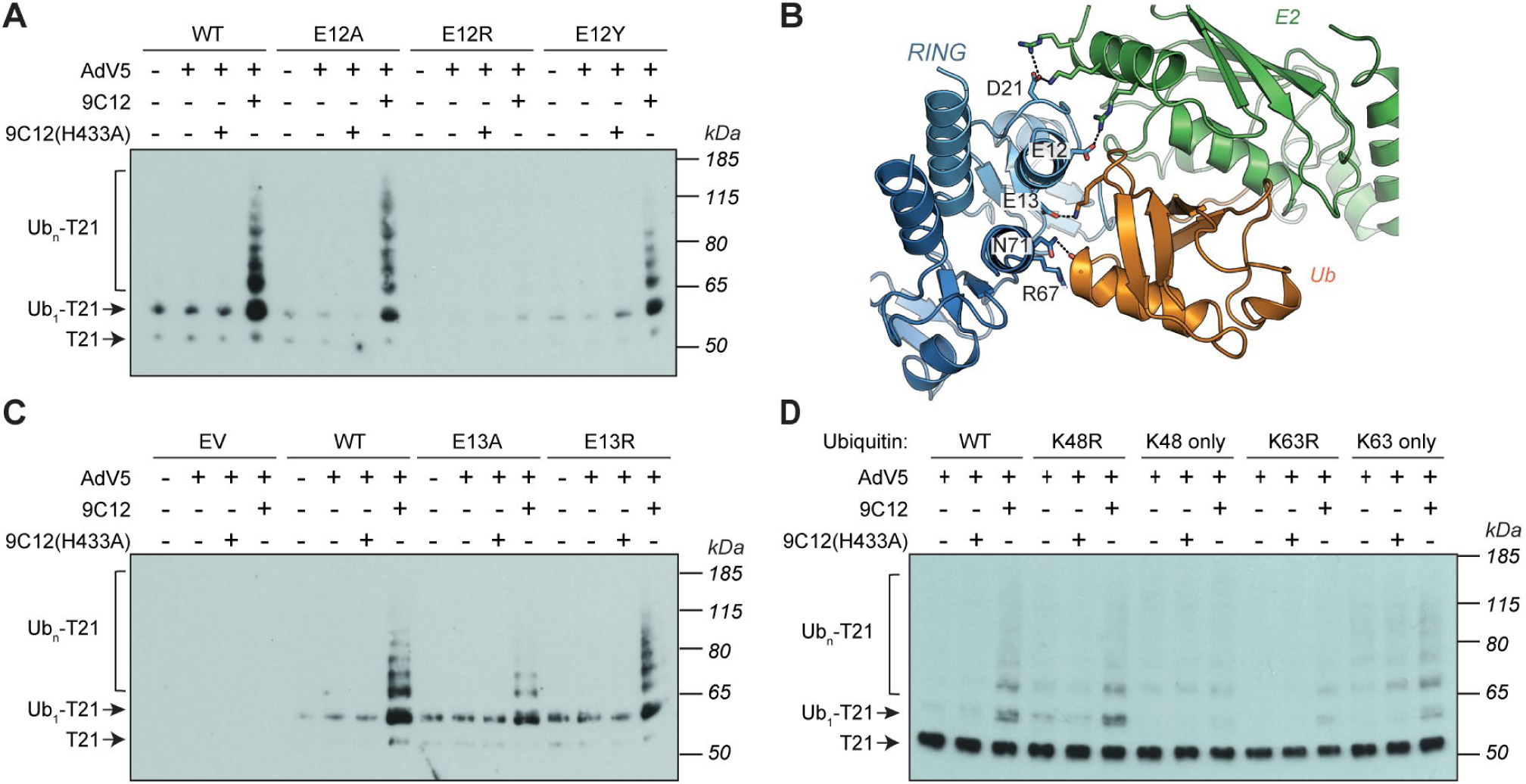
TRIM21 auto-ubiquitination upon substrate engagement. (A and C) HEK293T cells stably expressing the indicated TRIM21 mutants (EV = empty vector) were transfected with WT His-ubiquitin and infected with AdV5 ± 9C12 or 9C12(H433A). Ubiquitinated proteins were isolated by denaturing His-pulldown using Ni-NTA beads in 6M Guanidine buffer before immunoblot analysis with anti-TRIM21 antibody. (B) A co-crystal structure of TRIM21 RING:Ube2N∼ubiquitin complex (PDB: 6S53) showing the tri-anionic anchor motif (E12, E13 and D21) and second site residues (R67 and N71). (D) Immunoblot of TRIM21 following denaturing His-ubiquitin pulldown from HEK293T cells overexpressing the indicated ubiquitin mutant at 30 min post infection with AdV5 ± 9C12 or 9C12(H433A). See also Figure S1

### RING dimerization is required for TRIM21 activation

Next, we investigated how substrate binding might activate TRIM21 catalysis. TRIM21 is a dimeric multi-domain protein containing RING, B Box, Coiled-coil and PRYSPRY domains (Figure S2A). The Coiled-coil region of some TRIM family proteins can adopt an antiparallel conformation and it has been suggested to control activation by mediating oligomerisation (Esposito et al., 2017; Napolitano and Meroni, 2012). We found that the coiled-coil from TRIM21 undergoes monomer-dimer exchange but forms no higher order multimers (Figure S2B and C). Small angle x-ray scattering (SAXS) data (Figure S2D-F) revealed that TRIM21 exists as an elongated antiparallel coiled-coil with the two copies of the RING domain held apart by at least 150 Å (Figure 2A; METHODS). This arrangement did not change in the presence of IgG Fc (Figure 2B; Figure S2G-I; METHODS), suggesting that antibody binding does not induce large scale conformational changes in TRIM21.

**Figure 2.**
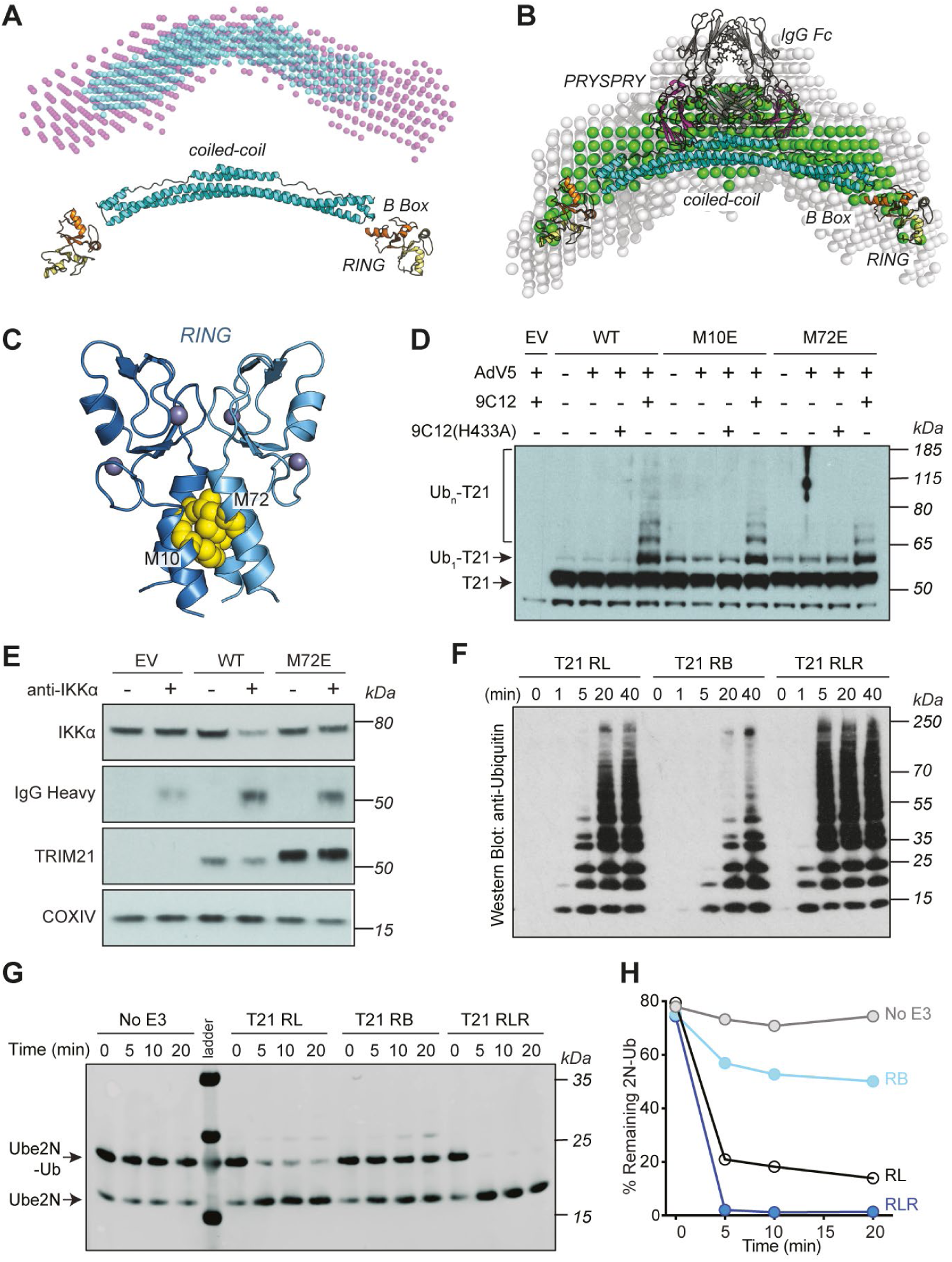
RING dimerization is required for TRIM21 activation. (A) Dummy atom model (DAM) structures of TRIM21 calculated from SAXS data shown in Figure S2D-F. Overlay of reconstructions of constructs comprising TRIM21 residues 1-235 (RBCC, purple spheres) and TRIM21 residues 129-235 (CC235, blue spheres). A structural model of the TRIM21 RBCC region based on atomic models of the TRIM21 RING (yellow) and B Box (orange) domains and TRIM25 coiled-coil domain (cyan) using PDBs 5OLM and 4CFG respectively. (B) SAXS-derived model of TRIM21:Fc complex. Averaged and filtered DAM’s (white and green spheres, respectively) overlaid with an atomic model of the TRIM21:Fc complex. The RING, B Box and coiled-coil regions are as above. The PRYSPRY (magenta) and IgG Fc (grey) domains are taken directly from PDB 2IWG. (C) Crystal structure of TRIM21 RING dimer showing the important residues M10 and M72 at the dimer interface. (D) Immunoblot of TRIM21 following denaturing His-ubiquitin pulldown from TRIM21 lentivector reconstituted HEK293T cells overexpressing WT His-ubiquitin at 30 min post infection with AdV5 ± 9C12 or 9C12(H433A). (E) Trim-away of IKKα in TRIM21 lentivector reconstituted HEK293T cell lines by electroporation of anti-IKKα IgG (anti-IKKα). Cell lysates were immunoblotted for the indicated proteins. (F) *in vitro* ubiquitination assay using the indicated TRIM21 constructs and Ube2N. Samples were taken at the indicated timepoints and analysed by immunoblotting with anti-ubiquitin antibody. (G) Immunoblot analysis of ubiquitin discharge from Ube2N by the indicated TRIM21 constructs over a course of 20 minutes. (H) Quantification of the Ube2N∼Ub band in G relative to time point 0 in each reaction. See also Figure S2

The large separation of the two RINGs within TRIM21, even when bound to IgG Fc, is significant because RING domains are typically only fully active as dimers (Buetow and Huang, 2016; Deshaies and Joazeiro, 2009). To investigate whether there is a functional requirement for TRIM21 RING dimerization we mutated two residues at the RING dimer interface (Figure 2C; (Dickson et al., 2018)). Introduction of an M72E mutation reduced TRIM21 polyubiquitin chain extension upon infection with Adv5+9C12, similar to that observed for the catalytic mutant E13A (Figures 1C and 2D). Moreover, this reduction in activity rendered TRIM21 unable to degrade substrate (IKKα) in a Trim-Away experiment (Figure 2E). Previously, it has been shown that monomeric TRIM21 RING possesses catalytic activity but is constitutively inhibited by the B Box (Dickson et al., 2018). However, comparing RING (RL), RING-B Box (RB) or RING-linker-RING (RLR; two RINGs in a single polypeptide) constructs revealed that enforcing RING dimerization significantly increases TRIM21 catalytic activity, as measured by polyubiquitin synthesis and ubiquitin discharge from Ube2N-Ub (Figure 2F-H). Thus, RING dimerization both increases TRIM21 catalysis *in vitro* and is necessary for TRIM21 activity in cells.

### Recruitment of multiple TRIM21 molecules to substrate induces RING dimerization

How can TRIM21 RING domains dimerise when they are held apart as monomers by an elongated coiled-coil? One potential mechanism for TRIM21 activation could be dimerization of RING domains between neighbouring TRIM21 molecules, for example on the surface of a virion (Figure 3A). This mechanism predicts that recruitment of multiple TRIM21 molecules to the virion is required for neutralisation. To test this, we took advantage of a neutralization assay in which the infectivity of adenoviral particles can be accurately quantified (Mallery et al., 2010). We varied ratios of WT to H433A anti-Adv5 hexon antibodies but maintained saturating concentrations, where around 200 antibodies have previously been shown to bind per virion (McEwan et al., 2012). Full levels of neutralization were abrogated when 9C12(H433A) reached >75%. By assuming WT and H433A antibodies were randomly distributed between viruses we found that ∼24 wildtype antibodies were required for neutralization (Figure 3B). Thus, efficient neutralization only occurs when multiple TRIM21 molecules are recruited to substrate. We further exploited the neutralisation assay to investigate whether the recruitment of multiple TRIM21 molecules switches on activity by facilitating RING dimerization. Introducing the dimerization mutant M72E significantly increased the antibody dose required for initiation of neutralization activity (Figure 3C). This lag phase in neutralization by the M72E mutant indicates that active RING dimers are formed upon recruitment of TRIM21 to antibody-coated virus. Detection of high levels of intracellular antibody-bound particles stimulates TRIM21 to activate NFkB signalling alongside is neutralization activity (Foss et al., 2016; McEwan et al., 2013). Accordingly, the M72E mutation also rendered TRIM21 unable to activate NFkB during infection, suggesting that the monomeric RING does not reach sufficient activity to overcome the high signalling threshold in these assays even at high antibody concentrations (Figure 3D). RING dimerization is required to allow efficient E2∼Ub recruitment; in a dimeric TRIM21 RING E3:E2∼Ub complex, the E2 contacts one RING while the ubiquitin contacts the second RING (Figure 1B; (Kiss et al., 2019)). To further test whether TRIM21 activation is driven by RING dimerization we introduced mutations N71D and R67A at the RING:Ub interface (Figure 1B). Both mutants behaved similarly to M72E and led to an increase in the critical threshold of antibodies required for neutralization (Figure 3E). These ubiquitin-binding mutations also rendered TRIM21 unable to activate NFkB during infection, decreased TRIM21 polyubquitination *in vitro*, and abolished TRIM21 activity during Trim-Away (Figure 3F-H). Altogether, these data suggest that substrate engagement drives TRIM21 RING dimerization, which induces TRIM21 ubiquitination activity.

**Figure 3.**
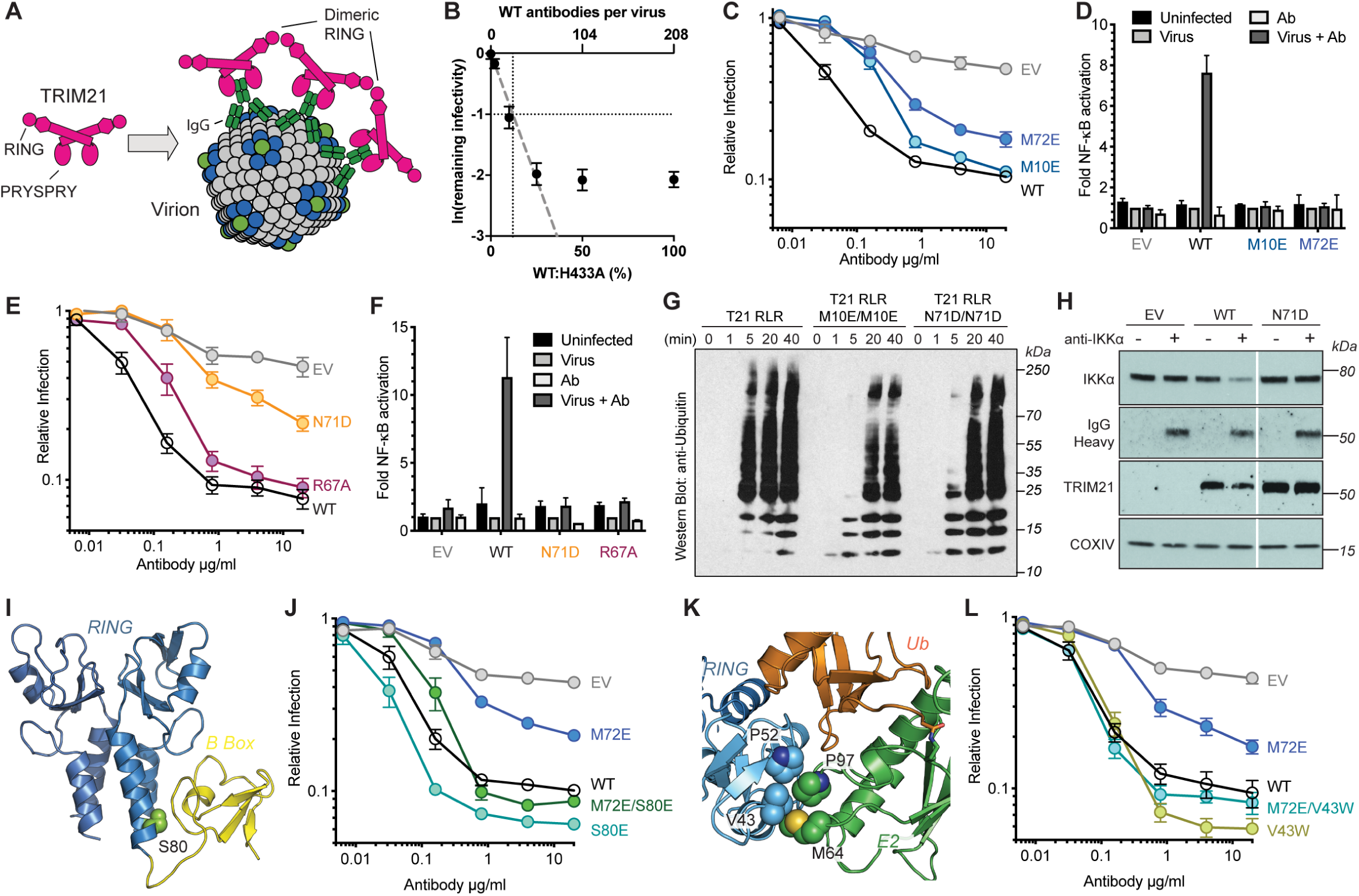
Substrate engagement induces RING dimerization and TRIM21 activation. (A) Schematic illustration of multimeric TRIM21 assembly on the surface of a virus-antibody complex. (B) Remaining infectivity of AdV5-GFP particles on HEK293T cells following incubation with saturating concentrations of 9C12. For 1/e neutralization, approximately 12% of antibodies must bear an intact TRIM21 binding site which is present on 9C12(WT) but lacking in 9C12(H433A). This equates to 24 out of the maximum ∼200 antibodies bound to AdV5 at saturation. (C, E, J and L) Neutralisation of AdV5 by 9C12 in lentivector reconstituted HEK293T cells expressing the indicated TRIM21 mutants. Data normalised to the virus only condition and presented as the mean ± SEM. (D and F) AdV5-9C12 immune complex-induced NF-kB activation in HEK293T cells stably expressing the indicated TRIM21 mutants. Data normalised to the virus only condition and presented as the mean ± SEM. (G) *In vitro* ubiquitination assay using the indicated TRIM21 RING-linker-RING (RLR) constructs and Ube2N. Samples were taken at the indicated timepoints and analysed by immunoblotting with anti-ubiquitin antibody. (H) Trim-away of IKKα in TRIM21 lentivector reconstituted HEK293T cell lines by electroporation of anti-IKKα IgG (anti-IKKα). Cell lysates were immunoblotted for the indicated proteins. (I) Crystal structure of TRIM21-RING-B-box (PDB: 5OLM) showing residue S80 at situated at the RING:B Box interface. (K) Structure of the hydrophobic core at the TRIM21 RING:Ube2N interface (PDB: 6S53). See also Figures S3 and S4

### RING dimerization relieves B Box inhibition

The clustering of TRIM21 molecules onto a viral capsid to allow RING dimerization and activate ubiquitination is reminiscent of the mechanism employed by the antiretroviral restriction factor TRIM5, which assembles into a hexagonal lattice on the surface of an infecting HIV capsid. The TRIM5 lattice is held together via essential interactions between neighbouring B Box domains, bringing RINGs from three different TRIM5 dimers into close proximity. The three RINGs then form interchangeable dimers that ubiquitinate the remaining RING monomer (Fletcher et al., 2018; Wagner et al., 2016). To test whether TRIM21 uses the same B Box-induced lattice mechanism, we mutated putative B Box:B Box interface residues in TRIM21 based on superposition of the TRIM21 B Box structure onto that of the trimeric TRIM5 B Box (Figure S3A). None of the mutations affected Adv5 neutralization or signalling (Figure S3B-D). To rule out the possibility that a different B Box:B Box interface is used to mediate higher order assembly, we also removed the B Box domain entirely. This ΔBox construct was still able to mediate efficient Adv5 neutralization (Figure S3E), suggesting that the TRIM21 B Box is not required to activate TRIM21 by assembling a lattice like TRIM5.

These results are consistent with published data showing that the TRIM21 B Box regulates catalytic activity by preventing E2∼Ub complexes from binding to the RING and driving the formation of active dimers (Dickson et al., 2018). We hypothesised that substrate-induced RING dimerization might be important to allow E2∼Ub to outcompete the B Box and overcome its autoinhibition (Figure S4A). To test this, we asked whether an S80E mutation, which displaces the B Box from the RING and relieves autoinhibition *in vitro* (Figure 3I; (Dickson et al., 2018)), could rescue activity of the dimerization-null mutant M72E (Figure S4B and C). Indeed, the S80E mutation restored the neutralization activity of the M72E mutant (Figure 3J). We sought to confirm the requirement for E2∼Ub to displace the B Box by asking whether M72E activity could also be rescued by improving E2 binding (Figure S4D). We introduced V43W, a structurally equivalent mutation to L51 in BRCA1, which increases E2 binding and is hyperactivating (Figure 3K; (Stewart et al., 2017)). Mutant V43W behaved analogously to S80E in neutralization experiments, increasing activity slightly when compared to wild-type and rescuing M72E (Figure 3L). Taken together, this suggests that RING dimerization is important to allow E2∼Ub to out-compete inhibitory B Box binding (Figure S4A-D).

### Trim-Away requires recruitment of multiple TRIM21 molecules to the target

The above data support a model in which TRIM21 becomes catalytically active when RING dimerization is induced by the clustering of multiple TRIM21 molecules on antibody-coated substrate (Figure 3A). We set about testing this model by using an artificial substrate containing a repeated epitope sequence. To this end, we designed a series of monomeric EGFP (mEGFP) reporter constructs with four copies of the 10 amino acid c-myc tag at the N-terminus. We serially introduced the point mutations L4A and I5A, which ablate binding by the monoclonal anti-myc antibody 9E10 (Hilpert et al., 2001). Critically, this antibody cannot induce antigen cross-linking owing to its atypical binding mode wherein both Fab domains contribute to antigen binding in a ‘clasping’ manner (Krauss et al., 2008). This resulted in a series of constructs with between zero and four anti-myc antibody binding sites per mEGFP (Figure 4A). Upon electroporation of anti-myc antibody, we observed efficient degradation of mEGFP and loss of fluorescence only in cells expressing mEGFP with 4 active anti-myc binding sites (Figure 4B-D). Degradation of the 4myc construct was antibody dose-dependent, whereas there was no loss of mEGFP containing only 3 anti-myc binding sites at any antibody concentration (Figure 4E). Finally, to confirm that antibody binding takes place to the 2myc construct, but that a critical number of antibodies has not been reached, we co-electroporated anti-IgG polyclonal antibody alongside anti-myc and observed efficient degradation (Figure 4F). These data confirm that TRIM21 activation requires the recruitment of multiple TRIM21 molecules to an antibody-coated substrate.

**Figure 4.**
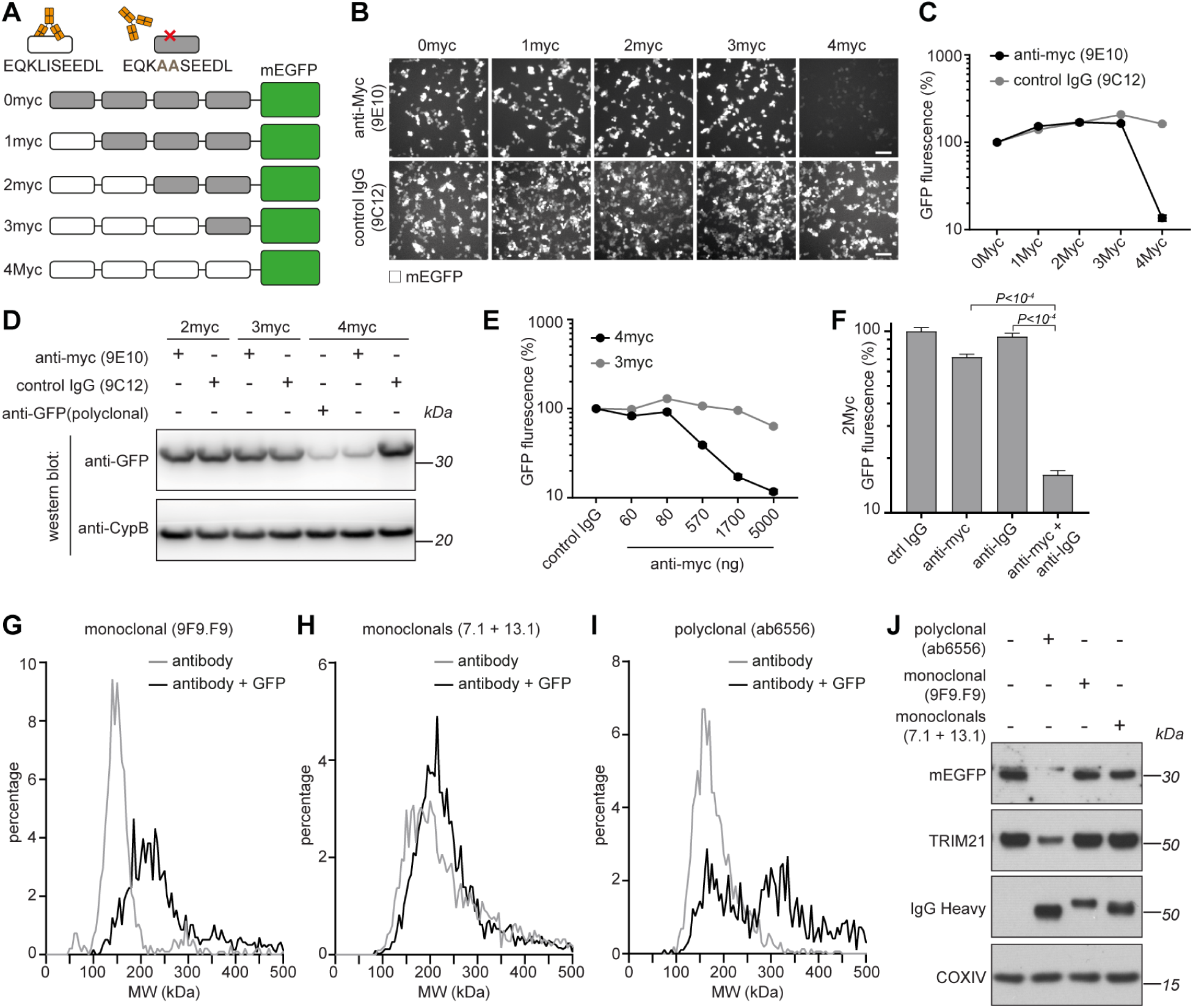
Trim-Away requires recruitment of multiple TRIM21 molecules to the target. (A) Schematic of myc-mEGFP constructs. (B-D) HEK293T-mCherry-TRIM21 cells were electroporated with mRNA encoding the indicated myc-mEGFP constructs together with either control IgG (9C12), ant-Myc (9E10) or anti-GFP (polyclonal) antibodies. 8 hours post-electroporation cellular GFP fluorescence was imaged (B) and quantified (C) using the IncuCyte system, or total GFP protein levels analysed by immunoblotting (D) cell extracts with the indicated antibodies. Scale bar 100 µm. (E) HEK293T-mCherry-TRIM21 cells were electroporated with mRNA encoding the indicated myc-mEGFP constructs together with control IgG or increasing concentrations of anti-Myc antibody and GFP fluorescence quantified 8 hours later. (F) HEK293T-mCherry-TRIM21 cells were electroporated with mRNA encoding 2myc-mEGFP together with the indicated antibodies and GFP fluorescence quantified 8 hours later. (G-I) The indicated antibodies (50 nM) either alone, or mixed with GFP protein (100 nM), were analysed by mass photometry. (J) RPE-1 cells expressing mEGFP were electroporated with the indicated antibodies and cell extracts blotted 3 hours later for the indicated proteins. See also Figure S5

We next used a system where we could directly quantify the number of antibodies bound to substrate. We mixed purified GFP together with either monoclonal or polyclonal anti-GFP antibodies and determined the molecular weight of the resulting protein complexes by mass photometry (Cole et al., 2017; Young et al., 2018). In the absence of GFP, all the antibodies used gave a peak of approximately 150kDa (Figure 4G-I, grey lines), consistent with the predicted size of a single IgG. For both a single monoclonal antibody (9F9.F9), and a mix of two monoclonals (7.1 + 13.1), the addition of GFP resulted in a peak shift to approximately 180-240 kDa, consistent with one or two GFP molecules (∼27kDa) bound by just a single antibody (Figure 4G and H, black lines). However, for the polyclonal antibody (ab6556), the addition of GFP resulted in the appearance of two peaks of approximately 180kDa and 330kD (Figure 4I, black line). This is consistent with a single GFP molecule bound by one (180kDa peak) or two (330kDa peak) antibodies. We did not observe larger molecular weight species, suggesting that a maximum of two antibodies could bind to GFP at any one time and that there was no antigen cross-linking leading to complexes containing more than two antibody molecules. We next tested which GFP:antibody complexes could trigger TRIM21 activation, and thus GFP degradation, inside cells. We electroporated cells expressing mEGFP with the anti-GFP antibodies. Strikingly, electroporation of polyclonal anti-GFP antibody (ab6556) resulted in efficient degradation of mEGFP, whereas both a single monoclonal (9F9.F9) and a mix of two monoclonal anti-GFP antibodies (7.1 + 13.1) did not (Figure 4J). TRIM21 was co-depleted together with mEGFP when using the polyclonal antibody (Figure 4J), consistent with TRIM21 activation only in this condition. All anti-GFP antibodies used were capable of binding GFP inside living cells, as shown by colocalization with membrane-anchored GFP following electroporation of antibodies into TRIM21 knockout cells (Figure S5). These data suggest that a threshold of two antibodies bound to substrate is required and can be sufficient to activate TRIM21.

### Molecular clustering is sufficient to activate TRIM21

In the above experiments, TRIM21 activation was induced by antibodies. However, if the only mechanistic requirement for activity is the clustering of TRIM21 molecules to facilitate RING dimerisation, then interaction with antibody should be unnecessary. To directly test this, we developed an experimental setup that would allow us to induce TRIM21 clustering independently of antibodies. To this end, we exploited the properties of engineered variants of the plant photoreceptor, Cryptochrome 2 (CRY2), which undergo clustering upon exposure to blue light (Bugaj et al., 2013; Más et al., 2000; Park et al., 2017; Taslimi et al., 2014b). We generated constructs with CRY2 fused to TRIM21 (Figure 5A) and compared their behaviour in human and *Drosophila* cells. Exposure of *Drosophila* S2 cells to blue light resulted in the rapid formation of cytoplasmic puncta for both mRFP-CRY2 and mRFP-CRY2-TRIM21 constructs, consistent with CRY2 clustering. Strikingly, live microscopy revealed that these clusters rapidly disappeared in cells expressing the TRIM21 fusion (mRFP-CRY2-TRIM21), but not in cells expressing mRFP-CRY2 alone (Figure 5B and C; Movie S1). The loss of CRY2-TRIM21 puncta was fully prevented by the addition of the proteasome inhibitor MG132, suggesting that this was due to proteasomal degradation. Moreover, these data indicate that TRIM21 clustering is sufficient to induce degradation even in insect cells that normally lack TRIM21. Western blotting confirmed that a TRIM21-CRY2 fusion protein expressed in human cells is also degraded via the proteasome following exposure to blue light (Figure 5D). The data demonstrate that molecular clustering is sufficient to activate TRIM21 independently of antibody binding.

**Figure 5.**
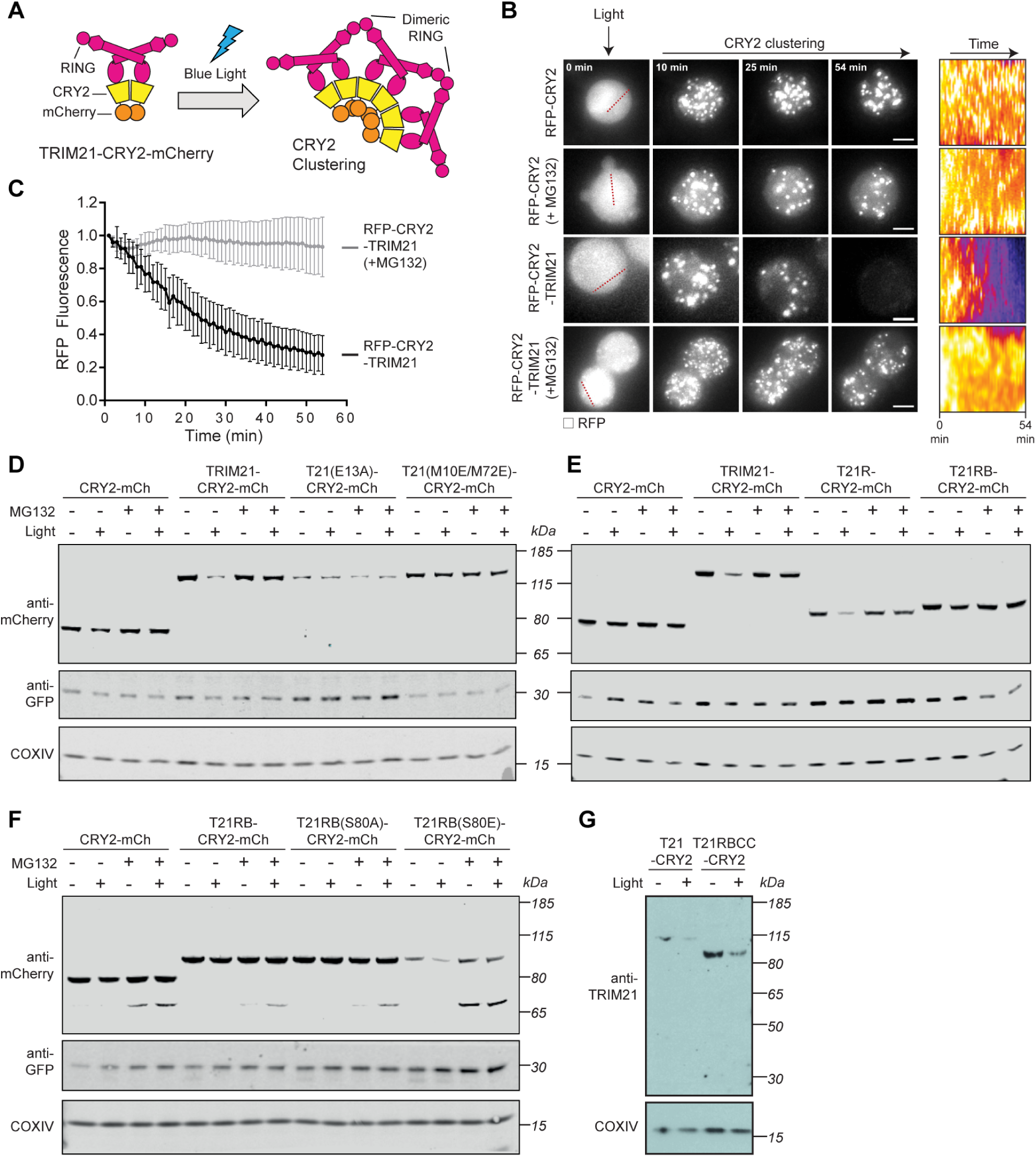
Molecular clustering activates TRIM21. (A) Schematic of light-induced clustering of TRIM21. (B-C) *Drosophila* S2 cells expressing the indicated constructs were incubated with or without MG132 and RFP fluorescence quantified by live imaging. Time shows minutes (min) from onset of blue light exposure. Scale bar 5 µm. Pseudo-coloured kymographs show fluorescence intensity in regions defined by red dotted lines. Graph shows mean fluorescence intensity (± SD) of RFP-CRY2-TRIM21 (n = 23) and RFP-CRY2-TRIM21+MG132 (n = 20) normalised for the respective controls (RFP-CRY2 (n=16) and RFP-CRY2 + MG132 (n=15)). (D-G) RPE-1 cells expressing the indicated constructs together with mem-mEGFP were incubated with or without MG132 and blue light for 3 hours prior to immunoblotting for the indicated proteins. See also Movie S2

Next, we used optogenetic clustering to investigate the mechanistic requirements for TRIM21 activation in human cells. First, we tested whether clustering-induced TRIM21 degradation requires its RING E3 ligase activity. Introducing the E13A point mutant, which prevents TRIM21 activation and autoubiquitination (Figure 1C; (Kiss et al., 2019)), was sufficient to prevent degradation of the TRIM21-CRY2 fusion upon light exposure (Figure 5D). Furthermore, the monomeric RING mutations M10E/M72E also prevented blue light-induced degradation of TRIM21-CRY2 (Figure 5D), suggesting that TRIM21 clustering specifically facilitates RING dimerization. Together this shows that TRIM21 is normally catalytically inert, but that upon molecular clustering active RING dimers form and drive degradation.

In a second series of experiments, we took advantage of light-induced clustering to ask which component domains are necessary and sufficient for TRIM21 activity. Surprisingly, we found that the minimum active construct was the RING domain alone, as the RING-CRY2 fusion protein (T21R-CRY2-mCh) was degraded to a similar extent as full length TRIM21-CRY2-mCh (Figure 5E). This result suggests that the coiled-coil and B Box domains of TRIM21 are required for regulation but not for activity. To investigate this further, we added the B Box to the active RING-CRY2 fusion (T21RB-CRY2-mCh) and found that degradation was now inhibited (Figure 5E). Degradation of the T21RB-CRY2 could be restored by introduction of an S80E mutation to displace the B Box and allow E2∼Ub binding but not by a control S80A mutation (Figure 5F). These experiments therefore support data from *in vitro* ubiquitination (Dickson et al., 2018) and Adv5 neutralization (Figure 3J) assays that the B Box is autoinhibitory. Interestingly, a CRY2-fusion protein containing the RING, B Box and coiled-coil (T21RBCC-CRY2) was efficiently degraded upon blue light exposure (Figure 5G). This suggests that the presence of the coiled-coil may assist in overcoming B Box autoinhibition. Consistent with this, addition of the coiled-coil (T21RBCC) could partially restore the *in vitro* ubiquitination activity of the TRIM21 RING-B Box (T21RB) construct (Figure S6A).

### Trim-Away without antibody Fc

With the discovery that TRIM21 activation and degradation can be uncoupled from antibody binding, we reasoned that it should be possible to engineer novel TRIM21 molecules that deliver targeted protein degradation independent of antibodies. To this end, we designed a TRIM21 chimera in which the PRYSPRY domain was replaced with an anti-GFP nanobody (Figure 6A; T21RBCC-vhhGFP4). We expressed this chimera, or the vhhGFP4-Fc fusion (Clift et al., 2017), in cells where GFP has been knocked-in to a single allele of Caveolin-1 (Shvets et al., 2015). As expected, expression of vhhGFP4-Fc, but not the TRIM21-null binding mutant vhhGFP4-Fc(H433A), caused efficient degradation of Caveolin-1-EGFP (Figure 6B). Remarkably, expression of the T21RBCC-vhhGFP4 chimera also led to Caveolin-1-EGFP degradation (Figure 6B). This suggests that TRIM21 can be directly recruited to, and degrade, target proteins independent of its Fc-binding activity.

**Figure 6.**
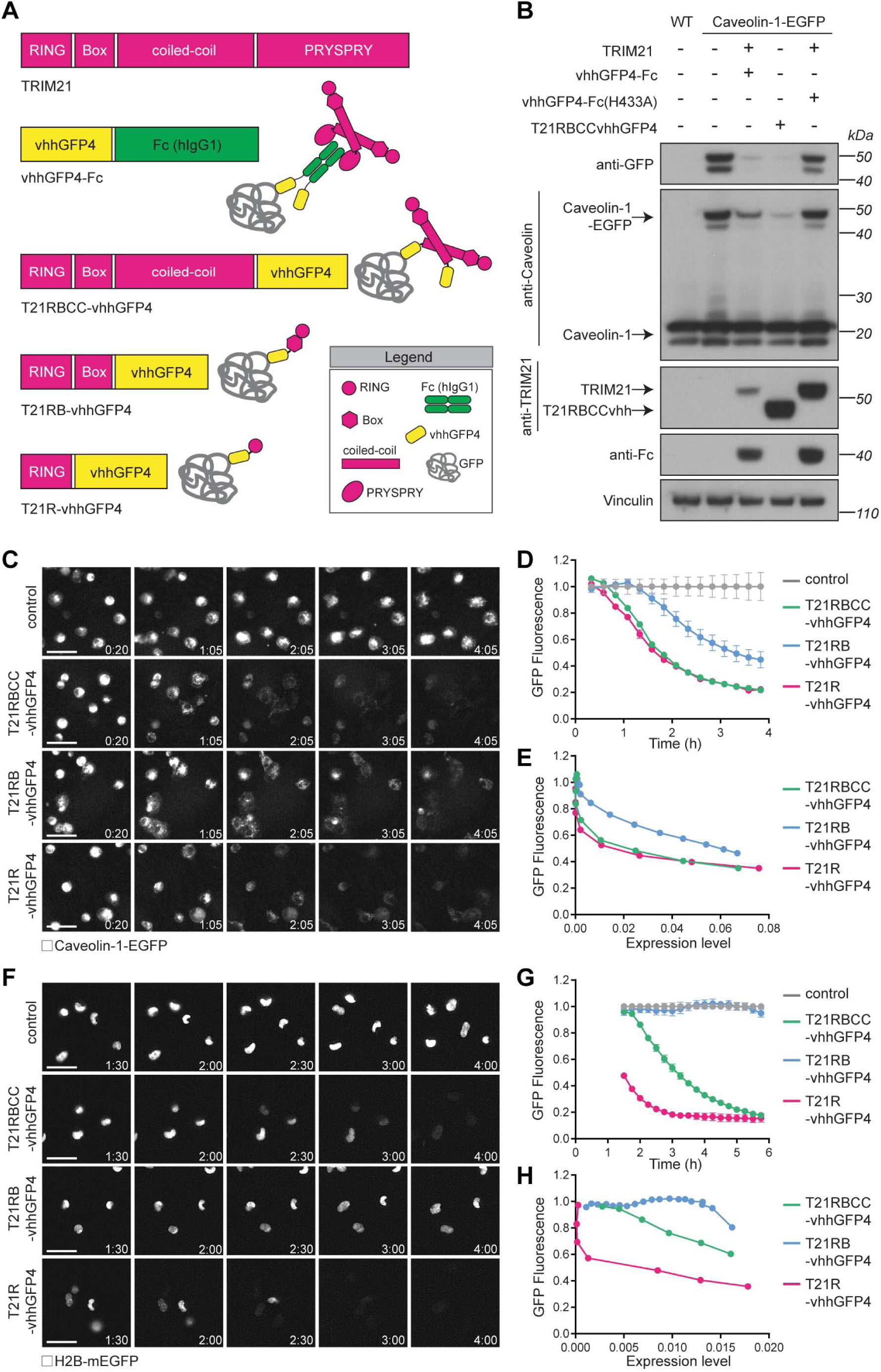
TRIM21-nanobody fusion proteins drive degradation. (A) Schematic of TRIM21, nanobody-Fc and TRIM21-nanobody chimeric constructs. (B) NIH3T3-Caveolin-1-GFP cells were electroporated with mRNA encoding the indicated constructs. 16 hours later cell extracts were immunoblotted for the indicated proteins. (C-D) NIH3T3-Caveolin-1-GFP and (F-G) RPE-1H2B-mEGFP-FKBP cells were electroporated with water (control) or mRNAs encoding the indicated TRIM21-nanobody chimeric constructs and GFP fluorescence was imaged (C and F) and quantified (D and G) using the IncuCyte system. Time shows hours:minutes (h:min) post-electroporation. Scale bar 50 µm (C) and 30 µm (F). (E) NIH3T3-Caveolin-1-GFP cells and (H) RPE-1-H2B-mEGFP-FKBP cells were electroporated with mRNA encoding mCherry-tagged versions of the indicated constructs and GFP and mCherry fluorescence quantified using the IncuCyte system. TRIM21-nanobody chimera expression levels (mCherry fluorescence) are plotted against GFP fluorescence. See also Figure S7 and Movie S2.

We expanded our TRIM21-vhh chimera approach to further engineer a minimal degrader molecule that combines targeting and catalytic activity in a single polypeptide. We designed a series of chimeric molecules containing different domains of TRIM21 fused to vhhGFP4 (Figure 6A). We induced expression of these molecules in either Caveolin-1-GFP knock-in cells, or cells stably expressing H2B-GFP, and monitored GFP degradation kinetics by live cell imaging. Strikingly, expression of a minimal construct containing only the TRIM21 RING domain fused to vhhGFP4 (T21R-vhhGFP4) caused rapid and efficient degradation of both Caveolin-1-GFP (Figure 6C and D; Movie S2) and H2B-GFP (Figure 6F and G). This was proteasome dependent, as treatment with MG132 prevented GFP degradation, despite colocalization of T21R-vhhGFP4 with Caveolin-1- and H2B-GFP (Figure S7A-D). Interesting, addition of the B Box domain to the RING (T21RB-vhhGFP4) caused a significant delay to the degradation of the GFP-tagged proteins, which could be rescued by further addition of the coiled-coil domain (T21RBCC-vhhGFP4; Figure 6A, C, D, F and G). The T21RB-vhhGFP4 chimera was expressed at lower levels compared to T21RBCC- and T21R-vhhGFP4 (Figure S7E and F). Nonetheless, plotting mCherry-TRIM21-vhh chimera expression levels against GFP fluorescence revealed that a higher concentration of T21RB-vhhGFP4 was required to induce GFP degradation (Figures 6E and H). Together these data are consistent with an inhibitory role for the B Box domain. Inhibition by the B Box was more noticeable when degrading H2B-GFP compared to Caveolin-1-GFP (Figure 6D and G). This may reflect the nature of the targets themselves, H2B is present in 2 copies within a histone octamer (Kanda et al., 1998), whereas Caveolin-1 is highly oligomeric (Hayer et al., 2010), which may help relieve B Box inhibition by allowing a high concentration of TRIM21 to assemble at caveolae, much like the surface of a virus. Importantly, these data show that a minimal RING-vhh construct (T21R-vhhGFP4) is the most efficient degrader, as it induced efficient degradation of target proteins even at low concentrations (Figures 6E and H). This is consistent with the enhanced activity of the RING domain compared to full length TRIM21 in *in-vitro* ubiquitination and ubiquitin discharge assays (Figure S6). Therefore, highly efficient protein degradation can be achieved by combining the catalytic activity of the TRIM21 RING together with substrate targeting in a single polypeptide chain.

### Expanding the Trim-Away toolbox

A key advantage of the original Trim-Away method is the direct delivery of protein reagents (e.g. antibody or antibody plus recombinant TRIM21) to the cell, which allows rapid target protein degradation even in primary cells that are sensitive to classical nucleotide delivery approaches (Clift et al., 2017). However, due to the size of whole antibodies (150kDa) acute targeting of nuclear-sequestered proteins is not possible with this approach. Furthermore, full length TRIM21 protein is difficult to produce in high quantities with high purity and activity (Clift et al., 2017; Clift et al., 2018). To overcome these problems, but maintain the advantages of protein delivery, we sought to generate a protein reagent based on the TRIM21-nanobody chimeras tested in Figure 6. We found that the T21R-vhhGFP4 protein expressed exceptionally well in bacteria and could be purified using a simple two-step protocol to give high yields and purity (Figure 7A; METHODS). Strikingly, electroporation of this protein into Caveolin-1-GFP knock-in cells resulted in dose-dependent degradation of Caveolin-1-GFP beginning immediately after protein delivery (Figure 7B). Addition of the B Box domain (T21RB-vhhGFP4) reduced the activity of the protein (Figure S7G-I), further supporting an inhibitory role for this domain. The small size of the T21R-vhh protein reagent (27kDa) makes it ideally suited to target proteins sequestered in the nucleus that cannot be accessed by whole antibodies. To test this, we electroporated the protein into primary mouse embryonic fibroblasts (MEFs) isolated from H2B-GFP transgenic mice. Remarkably, H2B-GFP was degraded within 1h of T21R-vhhGFP4 protein electroporation (Figure 7C). Protein delivery outperformed mRNA delivery in the primary MEFs (Figure 7C), likely because primary cells are sensitive to exogenous nucleotides, thus further highlighting the advantage of the T21R-vhh protein reagent for Trim-Away in primary cells.

**Figure 7.**
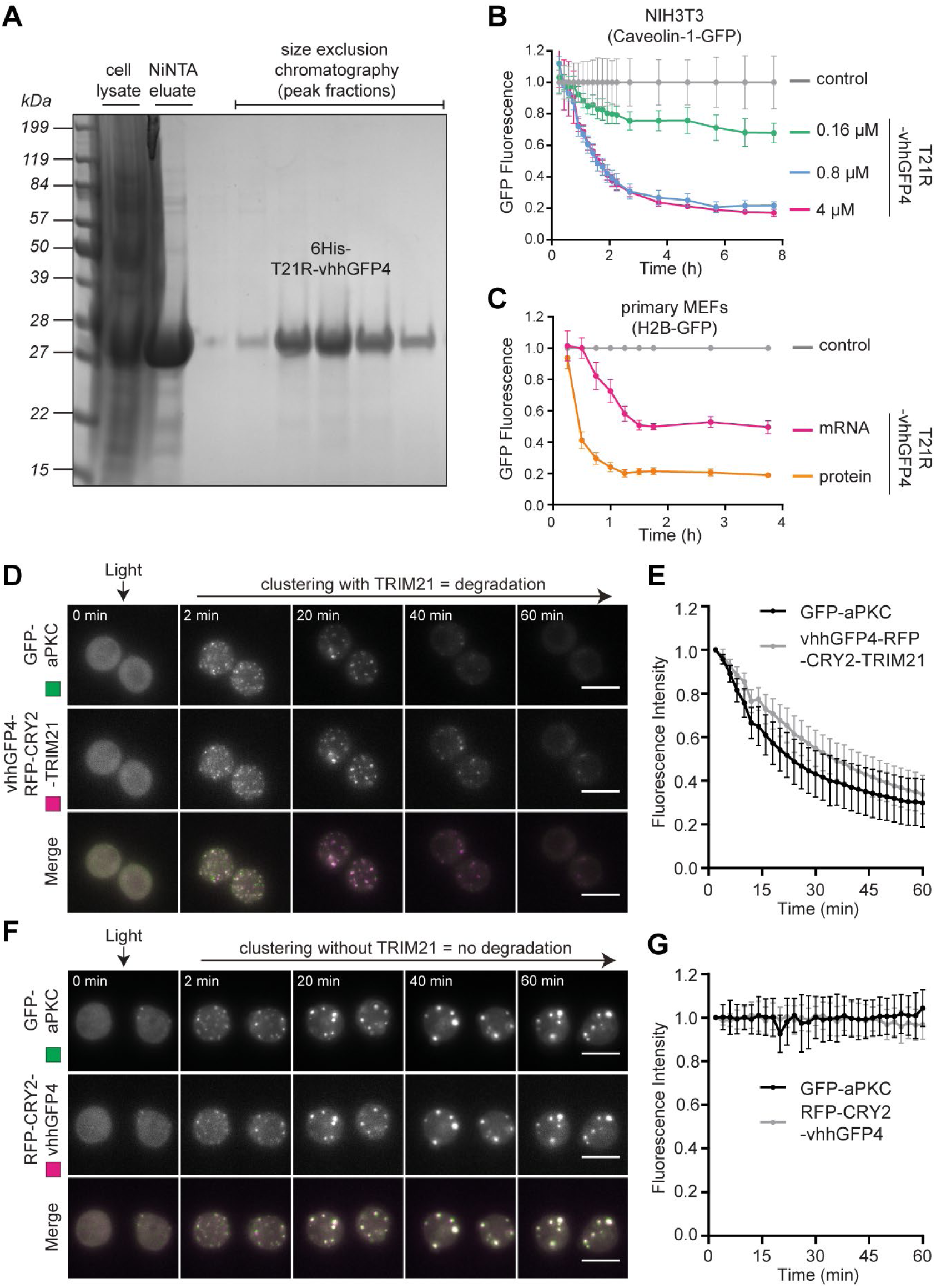
New ways to Trim-Away proteins. (A) Bacterially-expressed 6His-T21R-vhhGFP4 was purified using a two-step protocol of NiNTA-followed by size exclusion-chromatography and analysed by Coomassie blue staining of SDS-PAGE. (B) NIH3T3-Caveolin-1-GFP cells were electroporated with PBS (control) or the indicated concentrations of T21R-vhhGFP4 protein and GFP fluorescence was quantified using the IncuCyte system. (C) H2B-GFP primary MEFs were electroporated with PBS (control) or T21R-vhhGFP4 in the form or mRNA or protein and GFP fluorescence quantified using the IncuCyte system. Time shows hours (h) post-electroporation. (D-G) *Drosophila* S2 cells expressing GFP-aPKC and (D, E) vhhGFP4-RFP-CRY2-TRIM21 or (F, G) a LARIAT module that includes RFP-CRY2 fused with vhhGFP4 and which enables clustering in the absence of TRIM21. (E, G) Graphs show mean ± SD fluorescence intensity quantified by live imaging (n = 30 in E and n=24 in G). Time shows minutes (min) from onset of blue light exposure. Scale bar 10 µm.

We next asked if we could combine the TRIM21-nanobody chimeric approach with clustering-induced TRIM21 activation to achieve optogenetic control of Trim-Away. To this end, we generated a construct containing TRIM21 fused to both the GFP nanobody and light-controllable CRY2 domain (vhhGFP4-RFP-CRY2-TRIM21). We expressed this construct in *Drosophila* S2 cells together with an intended target of Trim-Away, GFP-tagged atypical protein kinase C (GFP-aPKC). At the onset of blue light stimulation, both proteins were distributed diffusely in the cytoplasm. However, within minutes of blue light stimulation GFP-aPKC formed clusters with RFP-CRY2-TRIM21 that were rapidly degraded (Figure 7D and E). Degradation of GFP-aPKC was TRIM21-dependent, because co-clustering of aPKC with RFP-CRY2 in the absence of TRIM21 [LARIAT construct; (Osswald et al., 2019)] did not lead to degradation (Figure 7F and G). These experiments were performed in *Drosophila* cells, which suggests this approach is applicable to model organisms where spatial control of Trim-Away may allow tissue-specific protein depletion. Taken together with the T21R-vhh protein approach, this works paves the way for both temporal and spatial control of Trim-Away in primary cells and model organisms.

## Discussion

Here we have investigated how the cytosolic antibody receptor and TRIM E3 ubiquitin ligase TRIM21 is activated and regulated. A significant finding of our study is that antibody binding is not a mechanistic requirement for TRIM21-mediated degradation. Antibodies do not induce a large-scale conformational change upon binding nor are they necessary to activate TRIM21 function in cells. Adenovirus infection assays and Trim-Away experiments with engineered targets demonstrate that degradation is induced by substrate-dependent recruitment of multiple TRIM21 molecules. Our data using a CRY2 light-inducible clustering system suggest that this is because molecular clustering activates TRIM21. The CRY2 data is also noteworthy because it demonstrates that TRIM21 achieves efficient degradation when it is both enzyme and substrate. This is consistent with published work (Fletcher et al., 2015), and data presented here, that TRIM21 modifies itself with a polyubiquitin chain. The combination of substrate-induced clustering and self-modification represents an alternative RING ubiquitination mechanism to the NEDD8-based activation system of Cullin RING Ligases (Baek et al., 2020). This different mechanism may explain how TRIM21 degrades diverse substrates; by ubiquitinating itself TRIM21 eliminates the requirement for a correctly orientated lysine residue within the antibody or substrate. This allows TRIM21 to target rapidly diverging pathogens during infection, and can be exploited to degrade any intracellular protein using Trim-Away technology.

Taken together, our data suggest the following mechanism of TRIM21 activation and regulation: TRIM21 is expressed as a catalytically inactive dimer, in which the two copies of its RING domain are located at either end of an elongated antiparallel coiled-coil. TRIM21 is kept inactive by two levels of regulation. First, separating the two RINGs in each TRIM21 molecule reduces activity with E2∼Ub, which requires a pre-formed RING dimer for efficient catalysis. Second, E2∼Ub is prevented from driving the equilibrium towards intermolecular RING dimerization by the B Box, which binds each RING monomer and occupies the E2 interface. Ubiquitination activity is activated by substrate-induced clustering of TRIM21 molecules, for instance by a viral particle with several antibodies bound, which promotes inter-molecular RING dimerization, making E2∼Ub binding energetically favourable and allowing B Box displacement. A key part of this mechanism is that the B Box only interacts with a RING monomer and therefore does not gain affinity upon dimerization, unlike E2∼Ub that interacts with both RINGs in the dimer. Once activated, TRIM21 assembles K63-linked ubiquitin chains on itself, which promotes degradation via the proteasome (Hofmann and Pickart, 2001; Ohtake et al., 2018; Saeki et al., 2009; Schrader et al., 2009), but can also induce immune signalling via TBK1 and IKK (McEwan et al., 2013). TRIM21 contains an IKK phosphorylation motif at the RING:Box interface (Dickson et al., 2018), suggesting there may be a feed-forward loop in which the RING is phosphorylated, favouring B Box displacement and enhancing E2∼Ub binding.

Our data also explain how Trim-Away works. TRIM21 is not activated by antibody binding alone. This means that antibodies delivered to the cytoplasm during Trim-Away are not immediately degraded by TRIM21, but instead are free to diffuse and reach their intended target. Only upon antibody-target engagement and subsequent clustering of TRIM21 molecules is TRIM21 activated to drive degradation. Indeed, TRIM21 and antibody have been shown to persist in the cytoplasm for many hours during zebrafish development before being rapidly degraded only upon target protein expression (Chen et al., 2019). The requirement for molecular clustering to activate TRIM21 also exposes a potential limitation of the Trim-Away method: monomeric proteins targeted with a monoclonal antibody cannot be degraded. However, we show that this limitation can be overcome simply by co-delivering a secondary antibody together with the primary targeting antibody, which allows multiple TRIM21 molecules to be recruited to monomeric targets to induce their rapid degradation. Our analysis of GFP:antibody complexes suggests that as little as two antibodies bound to a monomeric target can be sufficient to allow TRIM21-mediated degradation.

Another important finding within this work is that the TRIM21 RING domain alone is sufficient to drive degradation. We have exploited this discovery to engineer new Trim-Away chimerics that allow specific protein depletion without antibodies by fusing TRIM21 components with a substrate-targeting nanobody. Genetically encoding Trim-Away into a single polypeptide may be useful in biological systems where antibody delivery is not possible and paves the way for therapeutic delivery via gene therapy vectors *in vivo*. We have also shown that TRIM21 RING-nanobody chimerics can be directly delivered to cells in protein form to achieve rapid and acute degradation. This circumvents both the necessity for endogenous TRIM21 expression or any genetic manipulation and may be particularly useful in primary cells. Finally, we exploited the TRIM21 clustering activation mechanism to develop a variation of Trim-Away that can be induced by light. Together, this work greatly expands the Trim-Away technology toolbox beyond the use of standard antibodies and allows more flexibility when designing protein depletion experiments utilising TRIM21. In addition to Trim-Away there are a growing number of methods available for acutely controlling endogenous protein levels (Baudisch et al., 2018; Caussinus et al., 2012; Clift et al., 2017; Clift et al., 2018; Deng et al., 2020; Fulcher et al., 2016; Gross et al., 2016; Ibrahim et al., 2020; Ju Shin et al., 2015; Lim et al., 2020; Ludwicki et al., 2019; Marschall et al., 2014; Portnoff et al., 2014; Traub, 2019; Yamaguchi et al., 2019). However, our work here on TRIM21 provides the first mechanistic understanding of how an E3 ligase can be re-directed to degrade diverse non-canonical substrates. Further work will be crucial to allow researchers to make an informed choice of which E3 ligase to choose for targeted degradation of their protein of interest, either as a research tool or therapeutic.

As TRIM21 is simultaneously a member of the Fc receptor and TRIM RING E3 ligase families it is useful to consider the data presented here in both contexts. Cell surface Fc receptors (FcRs) are activated by binding to multivalent immune complexes. Cross-linked FcRs become substrates for phosphorylation at intracellular motifs, which in turn promotes downstream signalling responses (Ravetch and Bolland, 2001). The results here demonstrate that despite divergent evolutionary origins and functional mechanisms, the molecular cue for activation of classical FcRs and TRIM21 is identical: the clustering of receptors following binding to multivalent immune complexes.

Higher order assembly has previously been proposed to drive TRIM E3 ligase activity but with the assumption that a defined quaternary structure is formed, mediated by coiled-coil or B Box domains (Esposito et al., 2017). TRIM5, with its formation of a highly ordered lattice across the surface of the HIV capsid, is the exemplar of this model (Wagner et al., 2016). However, our data on TRIM21 suggest that neither the B Box nor the coiled-coil domain are required for its activity. Instead, the data show that TRIM21 E3 ligase activity and the generation of active RING dimers is activated by substrate-induced clustering. Intriguingly, substrate clustering has been shown to activate the Nedd4 family of HECT domain E3 ligases (Mund and Pelham, 2018). Thus, substrate-induced clustering may be a general mechanism of activation utilised across the ubiquitin ligase family.

## Author Contributions

Conceptualization, D.C. and L.C.J.; Methodology, J.Z., M.O., W.A.M., E.M.S., D.C. and L.C.J.; Investigation, J.Z., A.F.S., A.M., M.O., J.L., D.J., C.D., N.R., C.J., M.V. and D.C.; Writing – Original Draft, D.C. and L.C.J.; Writing – Review & Editing, J.Z., A.F.S., M.O., W.A.M., E.M.S., D.C. and L.C.J.; Supervision W.A.M., E.M.S., D.C. and L.C.J.; Funding Acquisition, W.A.M., E.M.S and L.C.J.

## Acknowledgements

We would like to thank Jan Terje Andersson for the 9C12 antibody variants and members of the James Lab for helpful discussions. J.Z. was supported by a PhD Studentship from the Rosetrees Trust and the Frank Edward Elmore Fund (University of Cambridge). M.O. is part of the GABBA PhD program from the University of Porto. W.M. is a Lister Institute Prize Fellow and is supported by a Sir Henry Dale Fellowship jointly funded by the Wellcome Trust and the Royal Society (Grant Number 206248/Z/17/Z) and supported by the UK Dementia Research Institute which receives its funding from UK DRI Ltd, funded by the UK Medical Research Council, Alzheimer’s Society and Alzheimer’s Research UK. E.M.S. acknowledges FCT - Fundação para a Ciência e a Tecnologia (contract CEECIND/00622/2017 and grant PTDC/BEX-BCM/0432/2014) and project Norte-01-0145-FEDER-000029 supported by NORTE 2020. This work was supported by the MRC (UK; U105181010) and a Wellcome Trust Investigator Award (200594/Z/16/Z).

---

**Figure S1.**
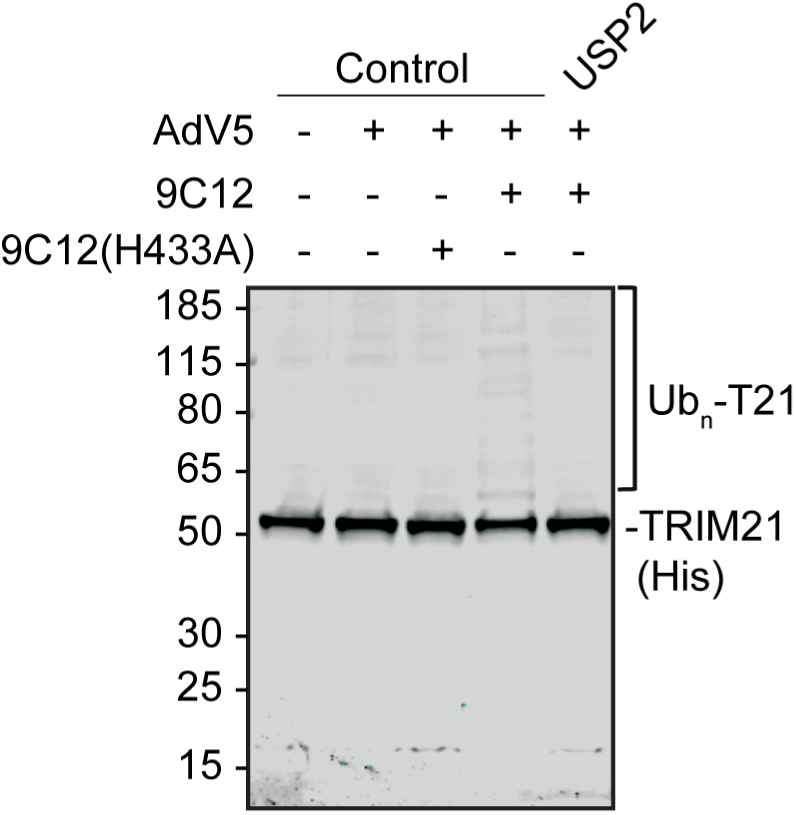
TRIM21 autoubiquitination with endogenous ubiquitin, related to Figure 1. Denaturing TRIM21-His pulldown from HEK293T cells stably expressing TRIM21-His and infected with AdV5 ± 9C12 or 9C12(H433A) which does not bind TRIM21. Cells were lysed in buffer containing 4M urea and where indicated, the Nickel beads were treated with the deubiquitinase USP2 before boiling and immunoblotting with anti-TRIM21 antibody.

**Figure S2.**
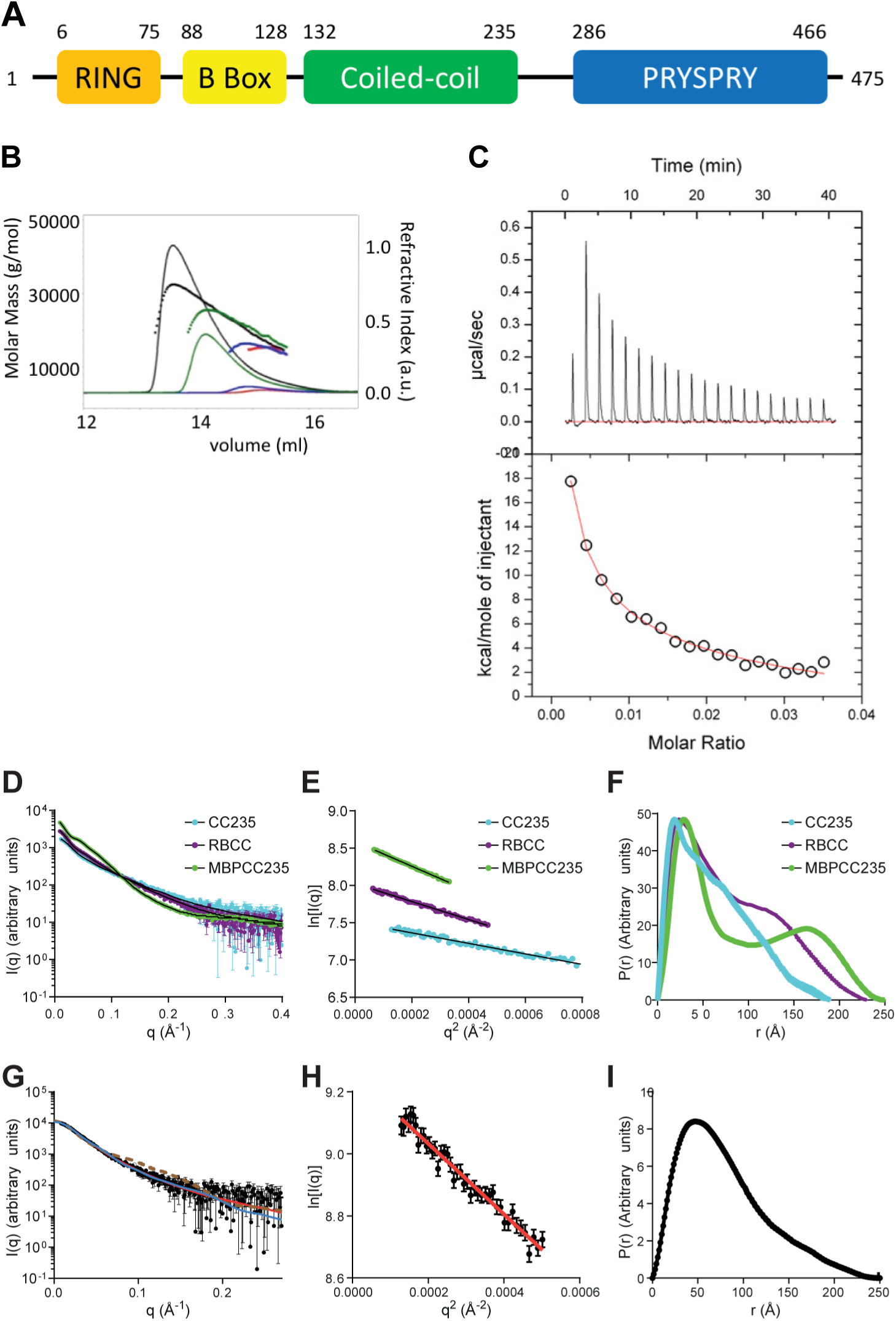
Structural and biophysical characterisation of TRIM21, related to Figure 2. (A) Sequence of TRIM21 with RING domain in gray, B Box in red, coiled-coil in cyan, L2 linker helices in orange & green and PRYSPRY in magenta. (B) SEC-MALS chromatograms of TRIM21 CC (129-235) loaded at concentrations of 15 (black), 5 (green), 0.55 (blue) and 0.25 mg/ml (red) are shown, in which the refractive index is indicated by the solid lines while the molar mass evaluated from the light scattering analysis is indicated with the corresponding coloured dotted lines. (C) Using a MicroCal iTC200 calorimeter, 140 µM TRIM21 coiled-coil was added in 2 µl injections into buffer at 25 °C. Integrated heats were then fit to a dimer dissociation model reveal an enthalpy of 65 kcal/mol and Kd of 7 µM. (D-F) SAXS data on TRIM21 constructs. (D) SAXS data were collected at the beam line P12 of the EMBL at the Petra-III storage ring (DESY, Hamburg). Protein expression, purification, data analysis and collection are in Supplementary Information. Data statistics are shown in Table 1. (D) Concentration-normalised scattering plots and DAM fits for CC235 (cyan, χ = 1.00), MBP-CC235 (green, χ = 1.31) and RBCC (purple, χ = 1.05). (E) The linear Guinier regions from (D). (F) Derived P(*r*) curves. The CC235 P(*r*) curve is consistent with an elongated rod, while the two peaks observed for MBP-CC235 and RBCC are consistent with ‘dumbbell’-shaped molecules. (G-I). SAXS data on TRIM21:Fc complex. (G) Scattering plot with DAM fit (red line, 1.00), TRIM21:Fc atomic model fit (blue, χ = 1.04), and apo-TRIM21 atomic model fit (brown dashed, χ = 2.00). (H) The linear Guinier regions from (G). (I) Derived P(*r*) curves.

**Figure S3.**
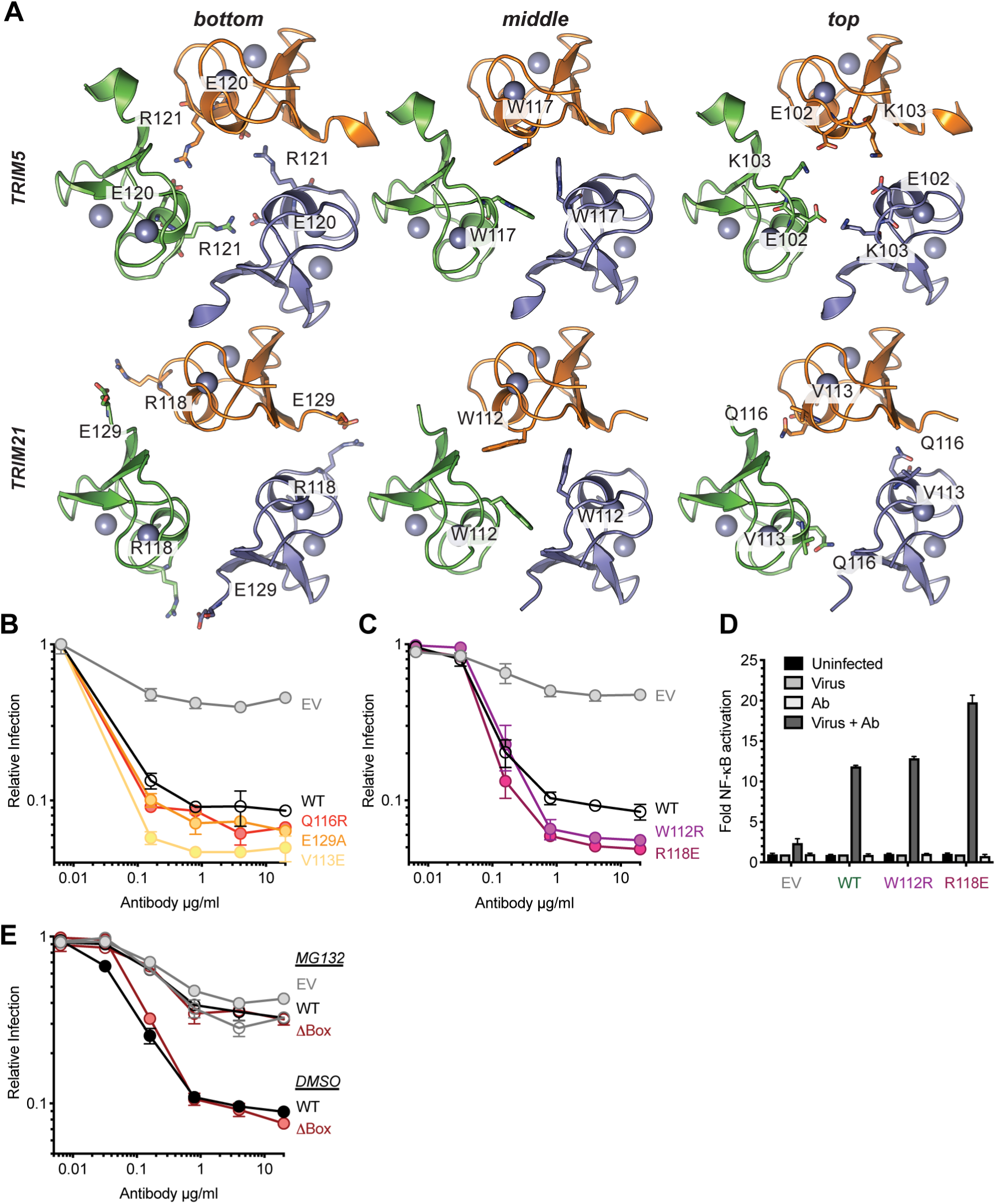
B Box domain is not required for TRIM21 activity, related to Figure 3. (A) Upper panel: TRIM5α (rhesus) residues involved in mediating the three layers of box-box interactions (PDB:5EIA). Lower panel: model of TRIM21 B box domain structure (PDB:5OLM) onto the trimeric box structure of TRIM5α (PDB:5EIA). Residues that could mediate B box: B box interactions are highlighted. (B and C) Neutralisation of AdV5 by 9C12 in lentivector-reconstituted HEK293T cells expressing the indicated TRIM21 mutants. Data normalised to the virus only condition and presented as the mean ± SEM. (D) AdV5-9C12 immune complex-induced NF-kB activation in HEK293T cells stably expressing the indicated TRIM21 mutants. Data normalised to the virus only condition and presented as the mean ± SD. (E) Neutralisation of AdV5 by 9C12 in HEK293T cells stably expressing TRIM21-ΔBox construct with or without MG132 (20 µM). Data normalised to the virus only condition and presented as the mean ± SD.

**Figure S4.**
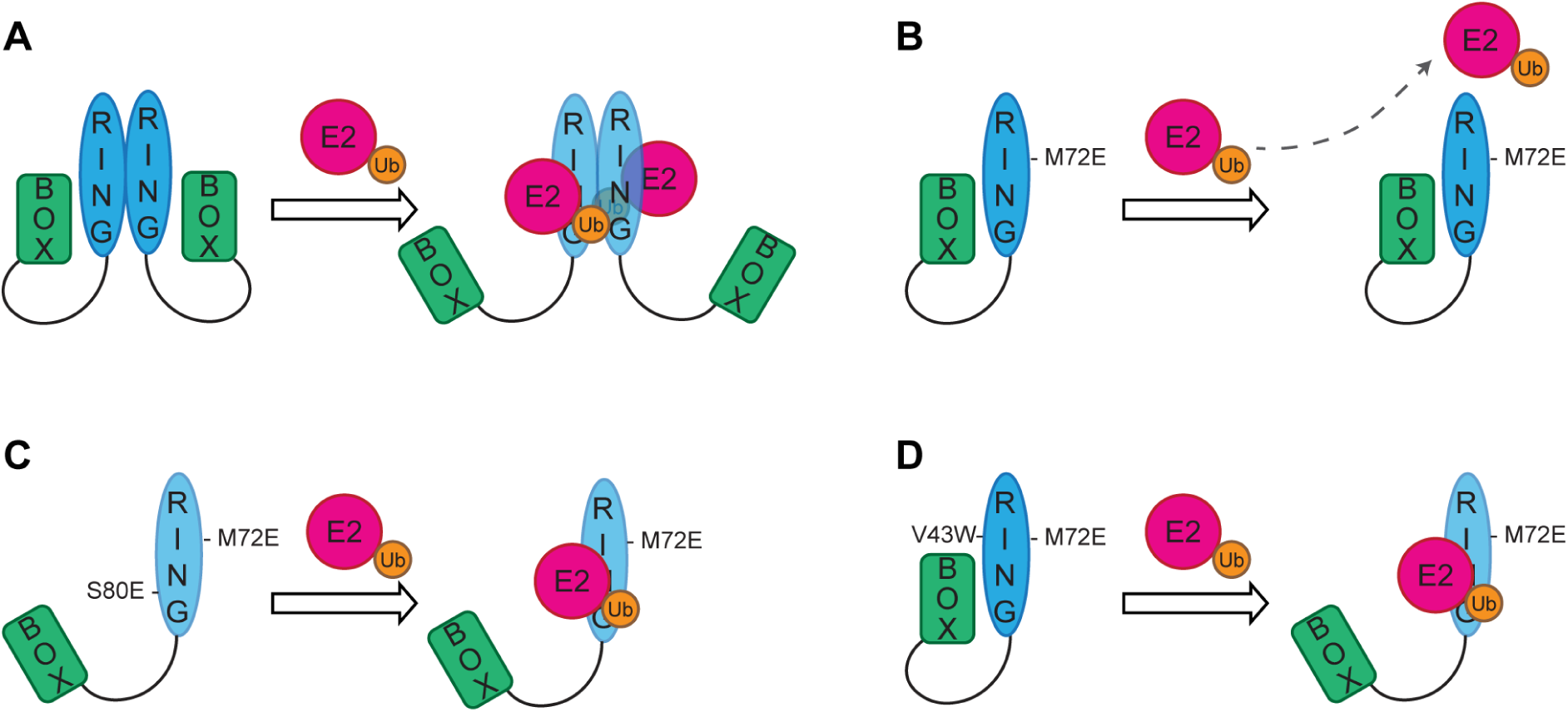
Models for RING domain regulation, related to Figures 2-3. (A) RING dimerization provides extra contacts between the charged ubiquitin (Ub) and the second RING protomer and perhaps lowering the activation energy required for the E2∼Ub to displace the autoinhibitory B Box. (B) RING dimerization mutants cannot dimerize, so it is more difficult for the E2∼Ub complex to displace the autoinhibitory B Box and bind to the monomeric RING domain. (C) Introduction of the S80E mutation at the RING: B Box interface results in electrostatic repulsion so that the B Box can be more readily displaced by the E2∼Ub complex leading to increased probability of interaction between the RING and E2∼Ub which partially compensates for the inability of the RING to dimerize. (D) Activity of the monomeric RING domain can be increased by enhancing its interaction with the E2 enzyme to compensate for the loss of additional contacts provided by RING dimerization and enables the E2∼Ub to compete off the autoinhibitory B Box.

**Figure S5.**
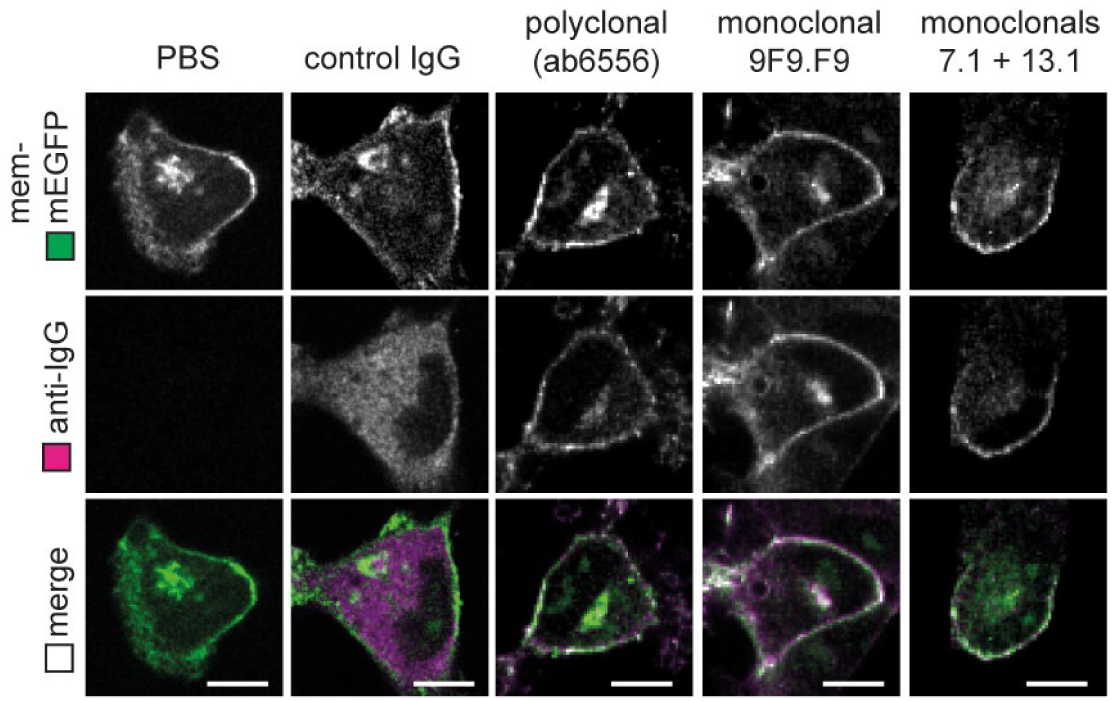
Anti-GFP antibodies bind to GFP inside living cells, related to Figure 4. RPE-1 TRIM21 KO cells expressing membrane-localised GFP (mem-mEGFP) were electroporated with PBS, control IgG or the indicated anti-GFP antibodies. Cells were fixed 3 hours post-electroporation and stained with alexa 647-conjugated anti-IgG secondary antibodies and imaged by confocal microscopy. Scale bar 10 µm.

**Figure S6.**
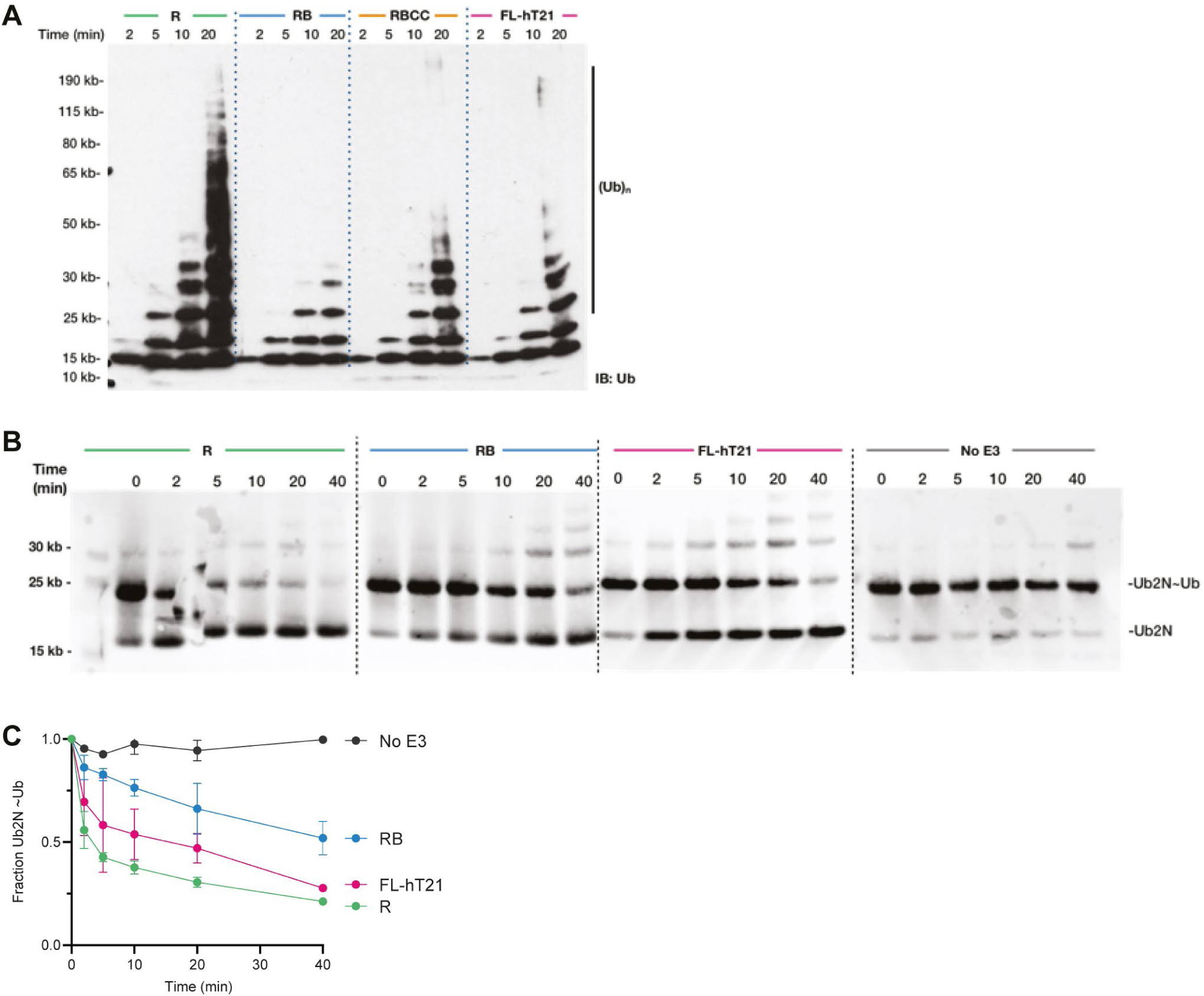
In vitro ubiquitination activity of TRIM21 constructs, related to Figures 5-6. (A) Catalysis of unanchored ubiquitin chains by TRIM21 RING (R), RING-Box (RB), RING-Box-Coiled-Coil (RBCC) and full length MBP-tagged TRIM21 (FL-hT21). (B-C) Catalysis of ubiquitin discharge from ubiquitin-conjugated Ube2N by RING (R), RING Box (RB), RING Box coiled-coiled (RBCC) and full length MBP-tagged TRIM21 (FL-hT21).

**Figure S7.**
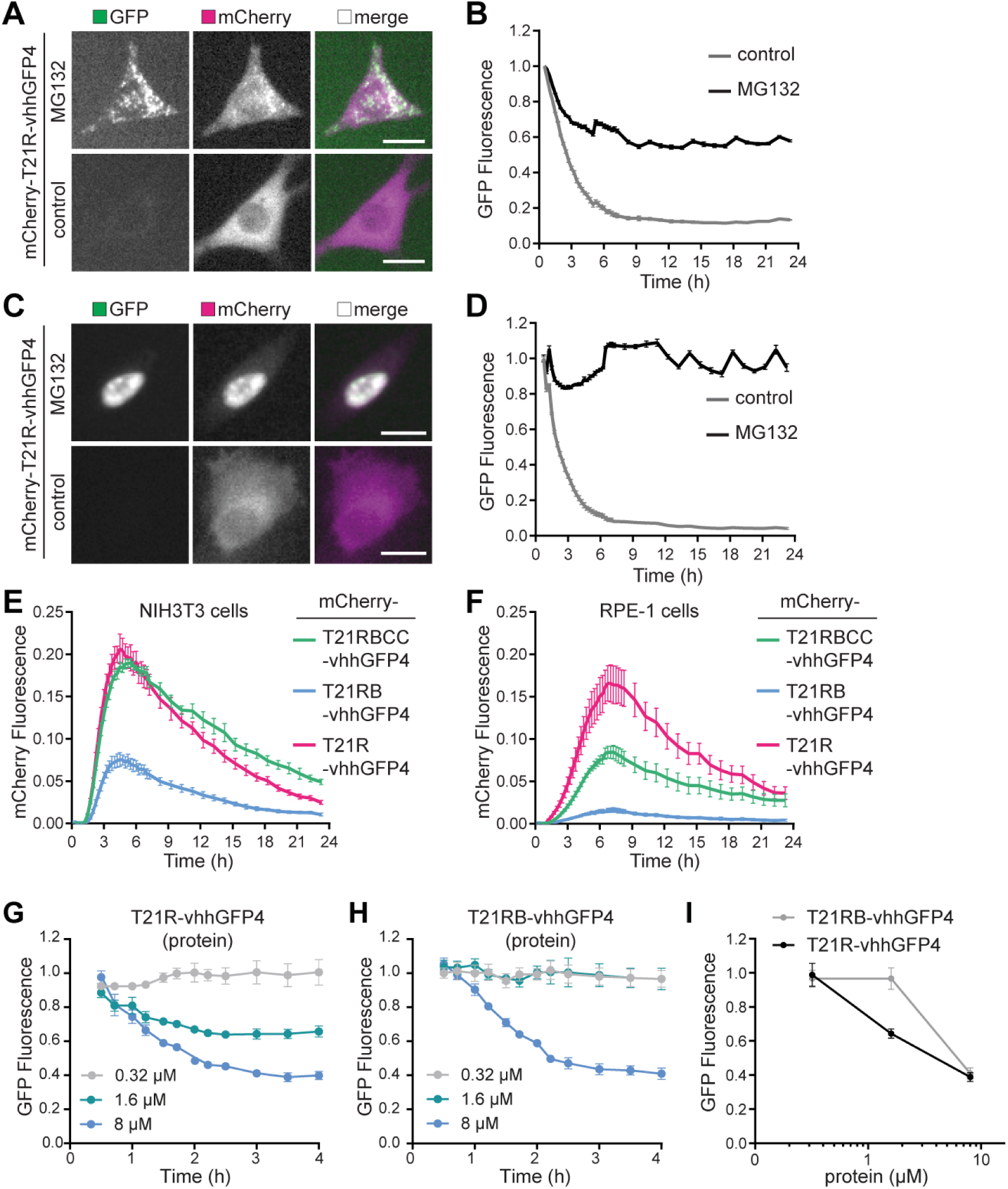
TRIM21-nanobody fusion proteins, related to Figure 6. (A-D) NIH3T3-Caveolin-1-GFP (A and B) and RPE-1-H2B-mEGFP-FKBP (C and D) cells were electroporated with mRNA encoding mCherry-T21R-vhhGFP4, incubated with either DMSO (control) or MG132 and imaged (A and C) and GFP fluorescence quantified (B and D) with the IncuCyte system. Scale bar 20 µm. (E and F) NIH3T3 cells (E) and RPE-1 cells (F) were electroporated with mRNA encoding mCherry-tagged versions of the indicated TRIM21-nanobody constructs and mCherry fluorescence quantified using the IncuCyte system. (G and H) NIH3T3-Caveolin-1-GFP cells were electroporated with the indicated concentrations of T21R-vhhGFP4 (G) or T21RB-vhhGFP4 (H) proteins and GFP fluorescence quantified with the IncuCyte system. (I) Data from G and H plotted as protein concentration against GFP fluorescence at 4 hours post-electroporation.

**Table S1.**
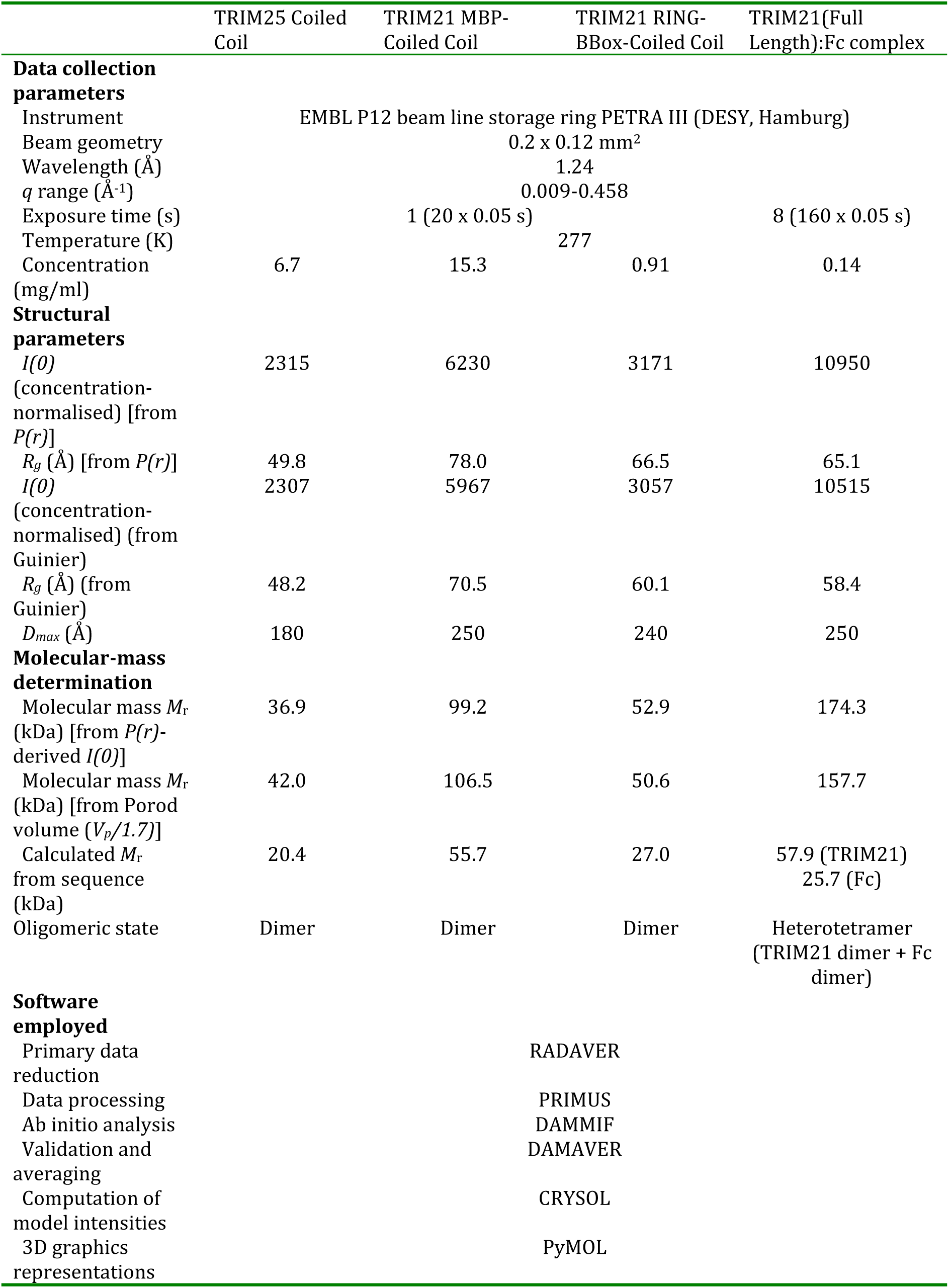
Crystallographic data collection, phasing and refinement statistics

## SUPPLEMENTAL MOVIE LEGENDS

**Movie S1. Light-induced clustering, related to Figure 5**

Time-lapse movie of *Drosophila* S2 cells expressing the indicated constructs showing that light-induced clustering of mRFP-CRY2clust-TRIM21 triggers protein degradation as fluorescence loss is inhibited by treatment with a proteasomal-inhibitor (MG132). Clustering was triggered by blue light at min 0. Scale bar = 5 μm.

**Movie S2. RING-nanobody fusion proteins, related to Figure 6**

Time-lapse movie of degradation of Caveoli-1-GFP with the indicated TRIM21-nanobody fusion proteins. Time shows h:min after electroporation of mRNA encoding the indicated TRIM21-nanobody constructs. Images obtained with the IncuCyte Live Cell Analysis system.

## METHODS

Further information and requests for resources and reagents should be directed to and will be fulfilled by the lead contact, Leo James.

### Mice

The experiments involving mice have been approved by the Medical Research Council Animal Welfare and Ethical Review Body and the UK Home Office under project licence number PCF3F9520. All mice were maintained in a specific pathogen-free environment according to UK Home Office regulations. H2B-GFP mice (CAG:H2B-EGFP) were obtained from The Jackson Laboratory (strain 006069).

### Mammalian Cell culture

HEK293T (ATCC) and NIH3T3-Caveolin-1-EGFP (Shvets et al., 2015) were cultured in DMEM medium (Gibco; 31966021) supplemented with 10% Calf Serum and penicillin-streptomycin. RPE-1 cells (ATCC) were cultured in DMEM/F-12 medium (Gibco; 10565018) supplemented with 10% Calf Serum and penicillin-streptomycin. All cells were grown at 37°C in a 5% CO_2_ humidified atmosphere and regularly checked to be mycoplasma-free. The sex of NIH3T3 cells is male. The sex of HEK293T and RPE-1 cells is female. Following electroporation, cells were grown in medium supplemented with 10% Calf Serum without antibiotics. For live imaging with the IncuCyte (Sartorius), cell culture medium was replaced with Fluorobrite (Gibco; A1896701) supplemented with 10% Calf Serum and GlutaMAX (Gibco; 35050061). For the RPE-1 cell CRY2 experiments, cells were changed into CO_2_ independent medium (Gibco; 18045088) supplemented with 10% Calf Serum and Glutamax (Gibco; 35050061). For the proteasome inhibition experiments MG132 (Sigma; C2211) was used at a final concentration of 25 µM.

### Drosophila S2 cell culture

Drosophila S2-DGRC cells (Drosophila Genomics Resource Center) were cultured at 25°C in Schneider’s Insect medium (Sigma-Aldrich, St. Louis, MO, USA) supplemented with 10% fetal bovine serum. Cells were transiently transfected using the Effectene Transfection Reagent (QIAGEN, Hilden, Germany), according to manufacturer’s instruction. Hsp70-promotor controlled constructs were induced by incubation at 37°C for 40 min at least 5h prior to cell imaging. After induction, cells were kept in the dark and manipulated under a 593nm LED light (Super Bright LEDs, St. Louis, MO, USA). For live imaging of S2 cells, cells were seeded on glass bottom-dishes (MatTek Corporation, Ashland, MA, USA) coated with poly-lysine and filmed in culture medium, at 25°C. All S2 cells were imaged between day 2 and day 5 post-transfection. For live proteasome inhibition assays, culture medium was supplemented with 20μM MG132 (Calbiochem) at least 1h before live imaging.

### Primary mouse embryonic fibroblasts (MEFs)

For MEF isolation, uteri isolated from 13.5-day-pregnant mice were washed with PBS. The head and visceral tissues were removed from isolated embryos. The remaining bodies were washed in fresh PBS, transferred into 3 ml of 0.25% trypsin, 1mM EDTA and minced using 2 scalpels. After 20 min incubation at 37°C cell suspension was mixed with 25ml of DMEM medium (Gibco; 31966021) supplemented with 10% Calf Serum and strained through 40 µm cell strainer. Cells were collected by centrifugation (200 × g for 5 min at 4°C), resuspended in fresh medium and seeded in T75 flask (passage 1). In this study, we used MEFs within three passages to avoid replicative senescence.

### Plasmids for lentivirus production

For stable expression of human TRIM21 in HEK293T cells, a native TRIM21 promoter-driven lentiviral vector (Zeng et al., 2019) was used. Point mutations and deletions within the TRIM21 coding sequence were generated using Quikchange mutagenesis (Agilent). For generation of the RPE-1-H2B-mEGFP-FKBP stable cell line the H2B-mEGFP (Addgene #105528) and FKBP (Promega N201A) coding sequences were inserted into a modified version of pSMPP (Addgene #104970) where the SFFV promotor and puromycin resistance sequences were replaced with PGK1 promoter and Zeocin resistance sequences respectively (pPMEZ) to generate pPMEZ-H2B-mEGFP-FKBP.

### Plasmids for transient transfection

For NF-κB signalling assays, plasmid pGL4.32 containing a firefly luciferase cassette under NF-κB response elements was obtained from Promega (Cat# E8491). For overexpression of ubiquitin, two copies of the 6His-ubiquitin sequence (Treier et al., 1994) were inserted in pCDNA3 (Invitrogen, V79020) to generate pcDNA3-2x-6His-Ub. Point mutations within the ubiquitin coding sequence were introduced using Quikchange mutagenesis (Agilent) to generate pcDNA3-2x-6His-Ub_K48R, pcDNA3-2x-6His-Ub_K63R, pcDNA3-2x-6His-Ub_K48-only (all lysine residues mutated to alanine except K48) and pcDNA3-2x-6His-Ub_K63-only (all lysine residues mutated to alanine except K63). To generate optogenetic constructs for S2 cell expression, vhhGFP4 (Saerens et al., 2005), mRFP (Campbell et al., 2002), mouse TRIM21 (Addgene #105516) and CRY2Clust (Park et al., 2017) coding sequences were inserted into heat-shock inducible Drosophila Gateway expression vectors (Life Technologies) to generate pHR-CRY2Clust-TRIM21, pHR-CRY2Clust and pH-vhhGFP4-RFP-CRY2Clust-TRIM21. The pHW-CIBN-MP_HRW-CRY2-V_H_H (Addgene #126024) and pHGW-aPKC plasmids have been previously described (Osswald et al., 2019).

### Plasmids for in vitro transcription of mRNA

To precisely control protein expression levels, constructs were expressed from in vitro transcribed mRNA that was electroporated into cells. This results in an approximately 24h window of expression where protein levels precisely correlate with the amount of mRNA electroporated. All constructs were clone into pGEMHE (Liman et al., 1992b), which contains UTR and polyA sequences for optimal mRNA stability and translation. pGEMHE-mEGFP and pGEMHE-membrane-mEGFP (Addgene #105526) were described previously (Clift et al., 2017). To generate the myc-mEGFP constructs, four copies of the c-myc epitope (EQKLISEEDL) with flexible linkers (GGGGS) were inserted upstream of mEGFP in pGEMHE-mEGFP to generate pGEMHE-4xmyc-mEGFP. Quikchange mutagenesis (Agilent) was used to generate the L4A/I5A mutations within the c-myc epitopes. To generate the light-inducible clustering constructs, the CRY2olig sequence (Taslimi et al., 2014a) was inserted with a c-terminal mCherry tag (Clontech) into pGEMHE to generate pGEMHE-CRY2olig-mCherry. TRIM21 mutants and truncations were inserted N-terminally to CRY2olig in pGEMHE-CRY2olig-mCherry with a flexible linker (GGGGS). The GFP nanobody-Fc fusion construct (pGEMHE-vhhGFP4-hIgG1-Fc; Addgene #105576) was described previously (Clift et al., 2017). Quikchange mutagenesis (Agilent) was used to generate the H433A mutation within the Fc region to prevent TRIM21 binding. To generate TRIM21-nanobody fusion constructs, TRIM21 sequences with or without an N-terminal mCherry tag were inserted upstream of the vhhGFP4 sequence with a flexible linker (GGGGS) into pGEMHE. For in vitro mRNA transcripton, pGEMHE plasmids were linearized and 5’-capped mRNA was synthesised with T7 polymerase (NEB HiScribeT7 ARCA kit) according to manufacturer’s instructions. For cellular expression from mRNA, typically 8 x 10^5^ cells (10.5 µl) were mixed with 2 µl of mRNA (∼0.5 µM) and electroporated using the Neon system. Typically, protein expression could be detected from 30 minutes post-electroporation and lasted ∼24 hours.

### Plasmids for protein purification

*E. coli* expression vectors encoding Ubiquitin, Ube1, Ube2N(K92R), T21R (amino acids 1-88) and T21RB (amino acids 1-129) have been described previously (Dickson et al., 2018; Fletcher et al., 2015; Kiss et al., 2019). To generate the TRIM21 RING-linker-RING construct (T21RLR), two copies of the TRIM21 RING domain separated by a GSH linker were inserted into an *E. coli* expression vector encoding an N-terminal 6His tag (pOPTH). Amino acids 1-285 of human TRIM21 (RING-Box-Coiled-coil) were inserted into an *E. coli* expression vector encoding an N-terminal GST tag (pOPTG) to generate pOPTG-T21RBCC. Amino acids 132-235 of human TRIM21 (Coiled-coil) were inserted into an *E. coli* expression vector encoding an N-terminal His tag (pOPTH) to generate pOPTH-T21CC (CC235). Amino acids 132-235 of human TRIM21 (Coiled-coil) were inserted into an *E. coli* expression vector encoding an N-terminal MBP tag (pOPTHM) to generate pOPTHM-T21CC (MBPCC235). To generate full length TRIM21 for protein production, MBP-tagged full-length TRIM21 was cloned into a pFastBac vector for expression in the baculovirus system (Bac-to-Bac, Invitrogen), resulting in pFastBac-MBP-TRIM21. To generate TRIM21-nanobody fusion constructs for protein production, TRIM21 sequences were inserted upstream of the vhhGFP4 sequence with a flexible linker (GGGGS) into pOPTH, resulting in the final constructs pOPTH-T21R-vhhGFP4 and pOPTH-T21RB-vhhGFP4.

### TRIM21 knockout cell lines

HEK293T TRIM21 knockout cells were described previously (Dickson et al., 2018). RPE-1 TRIM21 knockout cells were generated using the Alt-R CRISPR-Cas9 system from Integrated DNA technologies (IDT) with a custom-designed crRNA sequence (ATGCTCACAGGCTCCACGAA). Guide RNA in the form of crRNA-tracrRNA duplex was assembled with recombinant Cas9 protein (IDT #1081060) and electroporated into RPE-1 cells together with Alt-R Cas9 Electroporation Enhancer (IDT #1075915). Two days post-electroporation cells were plated one cell per well in 96 well plates and single cell clones screened by western blotting for TRIM21 protein. A single clone was chosen that contained no detectable TRIM21 protein and confirmed TRIM21 knockout phenotype in a Trim-Away assay.

### Stable cell lines

HEK293T-mCherry-hTRIM21 cells were described previously (Clift et al., 2017). Lentivirus particles were collected from HEK293T supernatant 3 days after co-transfection of lentiviral plasmid constructs together with HIV-1 GagPol expresser pcRV1 (a gift from Dr. Stuart Neil) and pMD2G, a gift from Didier Trono (Addgene plasmid #12259). Transfection was performed with Fugene 6 (Promega). Supernatant was filtered at 0.45 µm before storage at −80°C For stable expression of TRIM21, HEK293T TRIM21 KO cells (Dickson et al., 2018) were transduced with lentivirus particles at multiplicity ∼0.1 transducing units per cell and selected using puromycin at 2.5 µg/ml from 48 hours post-transduction. All TRIM21 constructs were expressed to similar levels as endogenous TRIM21 as confirmed by western blotting. For stable expression of H2B-mEGFP-FKBP, RPE-1 cells were transduced at multiplicity ∼0.1 transducing units per cell, sorted into single cells, and a single cell clone selected after screening for GFP expression by flow cytometry.

### Electroporation

Electroporation was performed using the Neon® Transfection System (Thermo Fisher). Cells were washed with PBS and resuspended in Buffer R (Thermo Fisher) at a concentration of 8 x 10^7^ cells ml^-1^. For each electroporation reaction 8 x 10^5^ cells (10.5 µl) were mixed with 2 µl of antibody or mRNA or protein to be delivered. The mixture was taken up into a 10 µl Neon® Pipette Tip (Thermo Fisher) and electroporated using the following settings: 1400V, 20 ms, 2 pulses. Electroporated cells were transferred to medium supplemented with 10% Calf Serum without antibiotics. Antibodies used for electroporation were rabbit anti-IκBα (E130, Abcam ab215972), mouse anti-myc (9E10; sigma, 05-419), mouse anti-Ad5 Hexon clone 9C12 (Foss et al., 2016), rabbit anti-mouse IgG (Jackson ImmunoResearch; 315-005-045), rabbit anti-GFP (Proteintech 50430-2-AP; 1:1000), rabbit anti-GFP (Abcam, ab6556), mouse anti-GFP (9F9.F9, abcam, ab1218), mouse anti-GFP (7.1+13.1, Roche, 11814460001) and normal rabbit IgG (Millipore 12-370).

### Ubiquitin and TRIM21 pull-down assays

Approximately 4 x 10^5^ HEK293T cells in a 6-well plate were transfected with 250 ng/well of pMT107-octa-6 His-ubiquitin or 200 ng/well of pcDNA3-2x-6His-ubiquitin constructs using 50 µl/well Opti-MEM (Gibco; 31985062) and 2 µl/well of FuGENE 6 Transfection reagent (Promega; E2691) following the manufacture’s protocol. Cell media were changed at 24 hours post-transfection. At 48 hours post-transfection, the media were replaced with 600 µl/well of serum-free CO_2_ independent medium (Gibco; 18045088) and the cells were chilled on ice for 30 min before infection with AdV5 or AdV5-9C12 complex. ΔE1ΔE3 AdV5-GFP (ViraQuest) was diluted 1:4 in PBS and mixed 1:1 with either 20 µg/ml of humanised 9C12 IgG3b or PBS and incubated for 1 hr at 20 °C to allow for complex formation. 48 µl of the virus or virus-antibody complex was added to each well in a 6-well plate with cold attachment on ice for 30 min. Following cold attachment, the cells were placed back into a 37 °C incubator for 30 min before the cells were washed twice with 1 ml/well of PBS and collected into clear 1.5 ml microtubes by scrapping. Cells were pelleted by centrifugation at 1000 x g for 5 min and the supernatants were removed before flash freezing the cell pellets in liquid nitrogen. For the denaturing His-Ubiquitin pulldowns, the cell pellets were lysed with 500 µl of buffer A containing 6M Guanidinium chloride, 5 mM imidazole 100 mM sodium phosphate buffer pH 7.4. The samples were mixed by vortexing and sonicated in ultrasonic water bath for 2 min (10s on/off) with amplitude to 40% to shear the DNA. 20 µl of buffer equilibrated Ni-NTA bead slurry (50% bead) were added to each sample and incubated for at least 3 hrs on a rotting wheel at 4°C. The Ni-NTA beads were pelleted by centrifugation at 16,000 x g for 30 s and washed two times with buffer A, two time with buffer B (buffer A and buffer C mixed 1:3) and two times with buffer C (20 mM Tris, 20 mM imidazole pH 6.8). Following the final wash, all of the supernatants were removed using 30 G needles and the beads were boiled for 5 min in 10 µl of 2 X LDS containing 100 mM DTT and 300 mM imidazole. The samples were then analysed by immunoblotting. For denaturing TRIM21-His pulldown, lysis buffer containing 4M Urea, 5 mM imidazole 50 mM Tris pH 7.4 was used and where indicated, the washed Ni-NTA beads were treated with USP2 (Boston Biochem; K-400) before immunoblotting.

### Adenovirus neutralisation assay

HEK293T Cells were plated at a density of 5 × 10^4^ cells/well in 0.5 ml of media in 24-well plates and were allowed to attach overnight. ΔE1ΔE3 AdV5-GFP (Viraquest) was diluted 1:1250 in PBS and mixed 1:1 with 9C12 antibody at the indicated concentrations and incubated for 1 hour at 20°C to allow for complex formation. For estimating the stoichiometry of TRIM21 for neutralization, 9C12(H433A) antibody was mixed at the indicated ratio with 9C12(WT) at 1 mg/ml before addition to AdV5-GFP particles. For the infection, 10 µl of the virus or virus-antibody complex was added to each well. Where indicated, 10 µM MG132 (Sigma) was added to the media 1 hour before infection. Cells were harvested by trypsinisation at 16-20 hrs post infection and evaluated for GFP expression on a BD LSRFortessa cell analyser (BD Biosciences). The results were analysed using FlowJo software (FlowJo LLC) and relative infection was calculated using the method described previously (Mallery et al., 2010).

### NFκB signalling assay

HEK293T cells were plated at a density of 3 x 10^5^ cells/well in a 6-well plate. On the following day, the cells were split 1:3 and left to incubate overnight. The next morning the cells were transfected with 200 ng/well of pGL4.32 NF-κB luciferase (Promega) using 50 µl/well Opti-MEM (Gibco; 31985062) and 2 µl/well of FuGENE 6 Transfection reagent (Promega; E2691) following the manufacture’s protocol. Cells were incubated for six-hours before reseeding at a density of 1 x 10^4^ per well in 96-well plates (Corning CellBIND; 3340) and allowed to attach overnight. ΔE1ΔE3 AdV5-GFP (ViraQuest) was diluted 1:3 in PBS and mixed 1:1 with either 20 µg/ml of humanised 9C12 IgG3b or PBS and incubated for 1 hr at 20 °C to allow for complex formation. 5 µl of the virus or virus-antibody complex was added to each well in triplicates and allowed to incubate for 6 hrs before the cells were lysed with 50 µl/well of steadylite plus luciferase reagent (Perkin Elmer; 6066751). The luciferase activity was quantified using a BMG PHERAstar FS plate reader.

### Light-induced CRY2 clustering

*Drosophila* S2 cells were transiently transfected with optogenetic constructs and between 2-5 days post-transfection expression induced by incubation at 37°C for 40 min at least 5h prior to cell imaging. After induction, cells were kept in the dark and manipulated under a 593nm LED light (Super Bright LEDs, St. Louis, MO, USA). The 470nm filter of a Nikon Ti motorized inverted epifluorescence widefield microscope (Nikon, Japan) was used to induce CRY2 clustering, whereas mRFP-Cry2clust modules were visualised using the 555 nm filter. For RPE-1 cells, following electroporation of mRNA, 2×10^5^ cells were plated in 24-well plates in antibiotics free media and allowed to adhere overnight. The following day, cells were media changed into CO_2_ independent medium with either DMSO or 20 µM MG132 before being exposed to blue light (wavelength of 450-500 nm) from a Dark Reader transilluminator (Clare Chemical). Cells were harvested for immunoblot analysis after 3 hours of exposure to blue light. The washed cell pellets were lysed directly in 60 µl of 1 x LDS sample buffer, sonicated in a water bath sonicator for 5 min (30s on/off, 4°C) and boiled at 95°C for 10 minutes before loading on to NuPAGE 4-12% Bis-Tris gels (Thermo Fisher).

### Microscopy

Live imaging of mammalian cells was performed using the IncuCyte S3 live cell analysis system (Sartorius) housed within a 37°C, 5% CO_2_ humidified incubator. Live imaging of *Drosophila* S2 cells was performed using a Nikon Ti motorized inverted epifluorescence widefield microscope (Nikon, Japan), equipped with a Lumencor SpectraX light engine and external emission filter wheel, and an Andor iXon888 camera (Andor, UK), with a PL APO LAMBDA 60X OIL/1,4 Oil WD 0,13mm objective, controlled by NIS elements 5.0 (Nikon, Japan) software. Z-stacks were acquired as 7 serial sections 1.5μm apart. The 470 nm filter was used to activate clustering and image GFP-aPKC, whereas mRFP-Cry2clust modules were imaged using a 555 nm filter. For fixed cells, images were acquired with a Leica SP8 microscope equipped with a 63x C-Apochromat 1.2 NA oil immersion objective.

### Measurement of fluorescence in live cells

To quantify RFP and GFP degradation in S2 cells, images were captured just before blue light exposure and in 1-minute (Figure 5) or 2-minute (Figure 7) intervals from blue light exposure onwards. Using ImageJ/FIJI software (Schindelin et al., 2012), the mean pixel intensity was measured within regions of interest (ROIs) that were manually tracked to include the complete area of each individual cell in which clustering occurred. Fluorescence intensity was quantified in average-intensity Z-projections covering all imaged planes and the signal was also measured in 3 non-transfected cells in the same sample, which were used for background subtraction. Fluorescence intensity over time is shown as the ratio between each timepoint and the second timepoint after light-induction. To quantify GFP and mCherry fluorescence in live mammalian cells, images were acquired and analysed using the IncuCyte live cell analysis system (Sartorius). Within the IncuCyte software, the integrated density (the product of the area and mean intensity) for GFP/mCherry fluorescence was normalised to total cell area (phase) for each image.

### Immunofluorescence

Cells were fixed for 30 min at 37°C in 100 mM HEPES (pH 7; titrated with KOH), 50 mM EGTA (pH 7; titrated with KOH), 2% formaldehyde (methanol free) and 0.2% Triton X-100. Fixed cells were incubated in PBS with 0.1% Triton X-100 overnight at 4°C. Antibody incubations were performed in PBS, 3% BSA and 0.1% Triton X-100. Antibodies used were Alexa Fluor 647-labelled anti-rabbit (Molecular Probes A21245; 1:400) and Alexa Fluor 647-labelled anti-mouse (Molecular Probes A21236; 1:400).

### Immunoblotting

Cells were washed in PBS, lysed in RIPA buffer (CST-9806) supplemented with a protease inhibitor cocktail (Roche), spun at 14000G for 10 min and cleared lysates mixed with NuPAGE LDS Sample Buffer and heated at 95°C for 5 mins. Samples were run on NuPAGE 4-12% Bis-Tris gels (Thermo Fisher) and transferred onto nitrocellulose membrane. Antibody incubations were mouse anti-ubiquitin (P4D1; Santa Cruz Biotehnology sc8017HRp; 1:1000), rabbit anti-Ube2N (Bio-Rad, AHP974, 1:1000), rabbit anti-IKKα (Y463, abcam ab169743; 1:5000), mouse anti-TRIM21 (D-12, Santa Cruz Biotechnology sc-25351; 1:500), rabbit anti-TRIM21 (D1O1D; CST-92043S; 1:1000), rabbit anti-COXIV (LI-COR 926-42214; 1:5000), rabbit anti-GFP (Proteintech 50430-2-AP; 1:1000), mouse anti-CyPB (k3E2; sc-130626; 1:1000), rabbit anti-mCherry (Abcam ab167453; 1:10000), mouse anti-GFP (7.1+13.1, Roche, 11814460001; 1:1000), rabbit anti-Caveolin (BD Biosciences 610059; 1:2000), goat anti-human IgG Fc (Biorad; 5211-8004; 1:2000) and rabbit anti-Vinculin (EPR8185; Abcam ab217171; 1:50,000). HRP-coupled secondary anti-mouse (Dako P0260), anti-mouse light chain specific (Millipore AP200P), anti-rabbit (Fisher 31462), anti-rabbit light chain specific (Millipore MAB201P) and anti-goat (Invitrogen A27014) were detected by enhanced chemiluminescence (Amersham, GE Healthcare) and X-ray films. IRDye-labelled secondary anti-Mouse IgG-IRDye 800CW (LI-COR; 925-32210; 1:5000) and anti-Rabbit IgG-IRDye® 680RD (LI-COR; 925-68071; 1:5000) were detected using LI-COR Odyssey CLx imaging system.

### Protein Purification

Ubiquitin, Ube1, Ube2N(K92R), T21R (amino acids 1-88) and T21RB (amino acids 1-129) were produced as described previously (Dickson et al., 2018; Fletcher et al., 2015; Kiss et al., 2019). The T21RLR and T21RBCC proteins were produced using the same protocol as T21RB (Dickson et al., 2018). To generate full length TRIM21 protein (T21FL), MBP-tagged TRIM21 (pFastBac-MBP-TRIM21) was expressed in SF9 insect cells in Lonza Insect-XPRESS according to standard protocols (Lonza). Cleared cell lysates were prepared by sonication of cell pellets in 50 mM Tris pH 8, 200 mM NaCl, 1 mM DTT, 10 μM ZnCl_2_, 15% (vol/vol) BugBuster (Novagen) and cOmplete protease inhibitors (Roche), followed by centrifugation 16,000 × g for 30 min. Lysates were loaded onto Amylose resin and the protein eluted with 10 mM Maltose. Eluted protein was passed through a HiLoad 16/600 Superdex 200 PG size exclusion column using an ÄKTA pure purification system (GE Healthcare). Peak fractions were pooled and concentrated using Amicon Ultra-4 centrifugal filter units (Millipore). Proteins MBP-T21CC (MBPCC235) and T21CC (CC235) were expressed in *Escherichia coli* C41(DE3) cells in 2TY medium and lysed as above. All proteins were purified by affinity (Ni^2+^- or Amylose-resin, as appropriate) and size exclusion chromatography (SEC) as above.

To generate TRIM21-nanobody fusion proteins, pOPTH-T21R-vhhGFP4 and pOPTH-T21RB-vhhGFP4 were transformed into *E. coli* C41 and TRIM21-nanobody protein expressed using autoinduction media (ZYP-5052) for 20 hours at 30 °C for T21R-vhhGFP4 and 4 hours at 37 °C followed by overnight expression at 17 °C for T21RB-vhGFP4. Cleared cell lysates were prepared by sonication of cell pellets in 50 mM Tris pH 8, 300 mM NaCl, 2 mM DTT, 10 mM Imidazole pH 8, 15% (vol/vol) BugBuster (Novagen) and cOmplete protease inhibitors (Roche), followed by centrifugation 16,000 × g for 30 min. Lysates were loaded onto Ni-NTA agarose (Qiagen), and protein eluted with 400 mM Imidazole. Eluted protein was passed through a HiLoad 10/300 Superdex 75 GL size exclusion column using an ÄKTA pure purification system (GE Healthcare). Peak fractions were pooled and concentrated using Amicon Ultra-4 centrifugal filter units (Millipore). For electroporation into cells, proteins were dialysed in PBS for 2 x 2 hours using Slide-A-Lyzer™ Dialysis Cassettes (Thermo Fisher). Protein aliquots were frozen and stored at −80°C until use.

### Ubiquitin discharge assay

Ube2N(K92R) was loaded with ubiquitin by mixing 40 µM of the E2 with 1 µM Ube1, 0.37 mM Ub and 3 mM ATP in 50 mM HEPES pH 7.5, 150 mM NaCl, 20 mM MgCl_2_, and incubating the reaction at 37°C for 30 min. The reaction was transferred to ice and used immediately. To observe E3 mediated discharge of ubiquitin, 2 µM ubiquitin loaded E2 was mixed with 1.5 µM TRIM21 protein constructs in 50 mM HEPES pH 7.5, 150 mM NaCl, 20 mM MgCl_2_, 50 mM L-lysine. Samples were taken at the time points indicated in the text and mixed immediately with LDS sample buffer at 4°C. The samples were boiled for exactly 20 s, resolved by SDS-PAGE and observed by immunoblot using rabbit anti-Ube2N (Bio-Rad, AHP974, 1:1000).

### Ubiquitin chain formation assay

Reactions were carried out in 50 mM Tris at pH 8, 2.5 mM MgCl2, 0.5 mM DTT with 0.2 mM Ub, 2 mM ATP, 1 µM Ube1, 0.5 µM Ube2N and 1.5 µM TRIM21 protein constructs. The reaction was started upon incubation at 37°C for the time points indicated in the text. The reaction was stopped via the addition of LDS sample buffer at 4°C, followed by boiling at 90°C for 2 min. The reactions were resolved by SDS-PAGE and ubiquitin detected by immunoblot using anti-ubiquitin-HRP (P4D1; Santa Cruz Biotechnology sc8017-HRP, 1:1000).

### Isothermal Titration Calorimetry (ITC)

ITC dilution experiments were performed using an iTC200 titration calorimeter (Malvern Panalytical). TRIM21 CC 235 was carefully dialysed into 20mM TRIS, pH 8, 100mM NaCl buffer at diluted to a concentration of 200uM. This was titrated into the same dialysis buffer in the ITC cell using twenty 2 uL injections. The resultant endothermic heat effects were integrated using Origin software supplied with the calorimeter.

### SEC-MALS

Size exclusion chromatography coupled to multi angle light scattering (SEC-MALS) measurements were performed using a Wyatt Heleos II light scattering instrument coupled to a Wyatt Optilab rEX online refractive index detector (Wyatt Technology). Samples of TRIM21 CC 235 (100uL) were resolved on a Superdex S-200 10/300 analytical gel filtration column (GE Healthcare) running at 0.5 ml/min in 20 mM Tris pH 8, 250mM NaCl buffer before passing through the light scattering and refractive index detectors in a standard SEC-MALS format. Protein concentration was determined from the excess differential refractive index based on 0.186 RI increment for 1 g/ml protein solution. The concentration and the observed scattered intensity at each point in chromatogram were used to calculate the molar mass from the intercept of the Debye plot using Zimm’s model as implemented in Wyatt’s ASTRA software.

### Small Angle X-ray Scattering (SAXS)

SAXS data (I(q) vs q, where q = (4πsinθ)/λ, where λ = 1.24 Å (10 kEV), 2θ = scattering angle), were collected at the beam line P12 of the EMBL at the Petra-III storage ring (DESY, Hamburg). Samples were collected at 277 Kelvin using an EMBL/ESRF new generation automated sample changer (1.5 mm sample capillary). Samples were measured while flowing through the capillary with scattering intensities recorded using 50 ms exposure times (20 frames total) on a 2D photon counting Pilatus 2M pixel X-ray detector (Dectris), maintaining a fixed sample to detector distance of 3.1 m (q-range 0.008-0.47 Å^-1^). The 2D-data (corrected for sample transmittance) were integrated and reduced using the automated software methods as described(Franke, 2012), to produce the final radially averaged, buffer subtracted 1D-scattering profiles. Based on comparison of successive frames, no detectable radiation damage was observed. All data manipulations were performed using the PRIMUS software package (Konarev et al., 2003).

The forward scattering at zero-angle I(0) and radius of gyration, *Rg*, were determined from Guinier analysis, assuming that at very small angles (*qRg* ≤ 1.3) the intensity is represented as *I(q)=I(0)*exp(-(*qRg*)^2^/3)). These parameters were also estimated from the full scattering curves using the indirect Fourier transform method implemented in the program GNOM(Svergun, 1992), along with the distance distribution function P(*r*) and the maximum particle dimensions D_max_. Molecular weights (MW) of solutes were estimated from SAXS data by comparing the extrapolated forward scattering with that of a reference solution of bovine serum albumin, and also from the hydrated-particle/Porod volume, *V_p_*, where MW is estimated as 0.625 times *V_p_*.

Linear Guinier regions and correct I(0)- and *V_p_*-derived MWs were used as quality control checkpoints prior to proceeding with further structural analysis (Jacques and Trewhella, 2010). For each sample, 12 *ab initio* dummy atom shape reconstructions were calculated using DAMMIF (Franke and Svergun, 2009) and averaged and filtered using DAMAVER. For the calculation of rigid-body model intensities, CRYSOL was used (Svergun et al., 1995).

To investigate the behaviour of the coiled-coil in solution, we collected SAXS data on a series of different constructs (Figure S2D-E, Table S1). The measured interatomic pair-distance distributions (P(*r*) curves; Figure S2F, cyan) for the coiled-coil suggest an elongated rod structure with a length of ∼190 Å and this was supported by the very large radius of gyration, *R_g_*, (46-52 Å) but small radius of gyration of cross-section, *R_c_* (7.5 Å). Adding an N-terminal maltose binding protein (MBP-CC235) as a ‘mass probe’ resulted in a clear bimodal P(*r*) curve, demonstrating that the coiled-coil must be an antiparallel dimer with the two MBP probe molecules at either end (Figure S2D-F, green). Replacing the MBP with the natural RING and B Box domains (RBCC), we observed a similar bimodal P(*r*) curve, albeit less pronounced given their significantly smaller mass (Figure S2D-F, purple). To compare the structures of the coiled-coil, MBP-CC235, and RBCC, we performed 12 dummy atom reconstructions using DAMMIF (typical fits shown in Figure S2D), which were then averaged and filtered using DAMAVER. An overlay of the Coiled-coil and RBCC *ab initio* structures is shown in Figure 2A and is consistent with a TRIM21 RBCC arrangement comprising an elongated antiparallel coiled-coil with the two copies of the RING domain held apart by at least 150 Å.

To test if antibody-binding causes large scale conformational change in TRIM21, we collected SAXS data on full-length TRIM21 protein in the presence of IgG Fc (Figure S2G-H, Table S1). The shape of the corresponding P(*r*) curve (Figure S2I) was no longer consistent with a dumbbell shape as with the RBCC but rather an elongated rod with a substantial mass towards the centre of the particle, suggesting that the PRYSPRY domains and Fc are positioned at the centre of the coiled-coil. To obtain a low resolution structure of the full-length TRIM21:IgG Fc complex, we performed twelve rounds of *ab initio* shape reconstruction (Figure S2H, red) to produce an averaged dummy atom model (Figure 2B, white) and a model for the most-populated volume (Figure 2B, green). A predicted rigid body model of TRIM21:Fc using published coordinates for component domains gave an excellent fit to the scattering data (χ = 1.0; Figure S2G, blue and 2B). In contrast, an ‘apo TRIM21’ model without bound Fc was a poor fit to the data (χ = 2.0, Figure S2G, brown). These data further support a structure in which the RING domains are located at either end of the coiled-coil with the PRYSPRY and bound Fc in the middle. Importantly, Fc binding did not induce any large-scale conformational changes in the coiled-coil and the RINGs remain monomeric. Consistent with this interpretation, the maximum linear dimension (*D_max_*) of the complex remains very similar that of the RBCC, while the curvature of the coiled-coil maintains a bend angle of 20° in all tested constructs irrespective of the presence of additional domains or Fc (Table S1).

### Mass Photometry

Refeyn One^MP^ was used to follow the complex formation of antibodies and GFP. In brief, solutions were prepared on ice and mixed reactions were incubated for at least 5 minutes before measurement. All experiments were prepared in 300 mM NaCl, 50 mM Tris pH 8 and 1 mM DTT. Mineral oil was used to mount a slide with a sample holder onto the objective. Immediately prior to measurement, 20 uL of sample had been applied onto the slide and the instrument focused according to manufacturer’s instructions. Data were processed and analysed using Discover MP v1.2.4 (Refeyn Ltd). Events were then normalised with a calibration curve and the resulting data exported and processed in GraphPad Prism.

### Statistical Analysis

Average (mean), standard deviation (s.d.), standard error of the meam (s.e.m) and statistical significance based on Student’s *t*-test (two-tailed) were calculated in Microsoft Excel or Graphpad Prism.

## KEY RESOURCES TABLE

| REAGENT or RESOURCE | SOURCE | IDENTIFIER |
| --- | --- | --- |
| **Antibodies** |  |  |
| Mouse anti-TRIM21 (D-12) | Santa Cruz Biotechnology | Cat#sc-25351; RRID:AB_628286 |
| Rabbit anti-TRIM21 (D1O1D) | Cell Signaling Technology | Cat#92043; RRID:AB_2800177 |
| Goat anti-adenovirus | Millipore | Cat#AB1056; RRID:AB_90213 |
| Mouse anti-Ad5 hexon (9C12) | (Foss et al., 2016) | DOI: 10.4049/jimmunol.1502601 |
| Human IgG1 anti-Ad5 hexon (9C12) | (Foss et al., 2016) | DOI: 10.4049/jimmunol.1502601 |
| Human IgG1(H433A) anti-Ad5 hexon (9C12) | (Foss et al., 2016) | DOI: 10.4049/jimmunol.1502601 |
| Mouse anti-ubiquitin (P4D1), HRP-conjugated | Santa Cruz Biotechnology | Cat#sc-8017HRP; RRID:AB_628423 |
| Rabbit anti-Ube2N (C-Terminal) | Bio-Rad | Cat#AHP974; RRID:AB_567248 |
| Rabbit anti-GFP | Abcam | Cat#ab6556; RRID:AB_305564 |
| Rabbit anti-GFP | Proteintech | Cat#50430-2-AP; RRID:AB_11042881 |
| Mouse anti-GFP (9F9.F9) | Abcam | Cat#ab1218; RRID:AB_298911 |
| Mouse anti-GFP (7.1+13.1) | Roche | Cat#11814460001; RRID:AB_390913 |
| Normal Rabbit IgG | Millipore | Cat#12-370; RRID:AB_145841 |
| Rabbit anti-mCherry | Abcam | Cat#ab167453; RRID:AB_2571870 |
| Mouse anti-myc (9E10) | Millipore | Cat#05-419; RRID:AB_309725 |
| Rabbit anti-mouse IgG (H+L) | Jackson ImmunoResearch | Cat#315-005-045; RRID:AB_2340038 |
| Rabbit anti-IKKα (Y463) | Abcam | Cat#ab59415; RRID:AB_733070 |
| Mouse anti-CyPB (k3E2) | Santa Cruz Biotechnology | Cat#sc-130626; RRID:AB_2169421 |
| Rabbit anti-Caveolin | BD Biosciences | Cat#610059; RRID:AB_397471 |
| Goat anti-human IgG Fc | Bio-Rad | Cat#5211-8004; RRID:AB_617100 |
| Rabbit anti-Vinculin (EPR8185) | Abcam | Cat#ab217171 |
| Rabbit anti-COXIV | LI-COR Biosciences | Cat#926-42214; RRID:AB_2783000 |
| Rabbit anti-Mouse IgG HRP-conjugated | Dako | Cat#P0260; RRID:AB_2636929 |
| Goat anti-Rabbit IgG (H+L) Cross-Adsorbed Secondary Antibody, HRP-conjugated | Thermo Fisher Scientific | Cat#31462; RRID:AB_228338 |
| Mouse anti-Rabbit light chain specific, HRP-conjugated | Millipore | Cat#MAB201P; RRID:AB_827270 |
| Goat anti-Mouse light chain specific, HRP-conjugated | Millipore | Cat#AP200P; RRID:AB_805324 |
| Rabbit anti-Goat IgG (H+L), Superclonal™ Recombinant Secondary Antibody, HRP | Thermo Fisher Scientific | Cat#A27014; RRID:AB_2536079 |
| Goat anti-human IgG (H+L)-HRP | Promega | Cat#W4031; RRID:AB_430835 |
| Goat anti-Mouse IgG, IRDye 800CW | LI-COR | Cat#925-32210; RRID:AB_2687825 |
| Goat anti-Rabbit IgG, IRDye 680RD | LI-COR | Cat#925-68071; RRID:AB_2721181 |
| Goat anti-Rabbit IgG (H+L)<br>Highly Cross-Adsorbed<br>Secondary Antibody, Alexa Fluor<br>647 | Thermo Fisher<br>Scientific | Cat#A21245; RRID:AB_2535813 |
| Goat anti-Mouse IgG (H+L)<br>Highly Cross-Adsorbed<br>Secondary Antibody, Alexa Fluor<br>647 | Thermo Fisher<br>Scientific | Cat#A21236; RRID:AB_2535805 |
| **Chemicals, Peptides, and Recombinant Proteins** |  |  |
| MG132 (Z-Leu-Leu-Leu-al) | Sigma | Cat#C2211 |
| Recombinant *A. victoria* GFP<br>protein | Abcam | Cat#ab84191 |
| USP2 | Boston Biochem | Cat#K-400 |
| Ubiquitin | (Dickson et al.,<br>2018) | DOI: 10.7554/eLife.32660 |
| Ube1 | (Dickson et al.,<br>2018) | DOI: 10.7554/eLife.32660 |
| Ube2N(K92R) | (Dickson et al.,<br>2018) | DOI: 10.7554/eLife.32660 |
| T21R | (Dickson et al.,<br>2018) | DOI: 10.7554/eLife.32660 |
| T21RB | (Dickson et al.,<br>2018) | DOI: 10.7554/eLife.32660 |
| T21RLR | This paper | N/A |
| T21RBCC | This paper | N/A |
| MBP-T21FL | This paper | N/A |
| MBP-T21CC (MBPCC235) | This paper | N/A |
| T21CC (CC235) | This paper | N/A |
| Human plasma IgG-Fc | Athens Research &<br>Technology | Cat#16-16-090707-FC |
| 6His-T21R-vhhGFP4 | This paper | N/A |
| 6His-T21RB-vhhGFP4 | This paper | N/A |
| **Critical Commercial Assays** |  |  |
| Neon transfection system | Thermo Fisher<br>Scientific | Cat#MPK5000S |
| HiScribeT7 ARCA kit | New England<br>Biolabs | Cat#E2065S |
| IncuCyte S3 live cell analysis<br>system | Sartorius | N/A |
| **Experimental Models: Cell Lines** |  |  |
| Human: HEK293T | ATCC | Cat#CRL-1573; RRID:CVCL_0045 |
| Human: HEK293T TRIM21 KO | (Dickson et al.,<br>2018) | DOI: 10.7554/eLife.32660 |
| Human: HEK293T-TRIM21_E12A | (Kiss et al., 2019) | DOI: doi.org/10.1038/s41467-019-12388-y |
| Human: HEK293T-TRIM21_E12R | (Kiss et al., 2019) | DOI: doi.org/10.1038/s41467-019-12388-y |
| Human: HEK293T-TRIM21_E12Y | (Kiss et al., 2019) | DOI: doi.org/10.1038/s41467-019-12388-y |
| Human: HEK293T-TRIM21_E13A | (Kiss et al., 2019) | DOI: doi.org/10.1038/s41467-019-12388-y |
| Human: HEK293T-TRIM21_E13R | (Kiss et al., 2019) | DOI: doi.org/10.1038/s41467-019-12388-y |
| Human: HEK293T-TRIM21_M10E | This paper | N/A |
| Human: HEK293T-TRIM21_M72E | This paper | N/A |
| Human: HEK293T-TRIM21_R67A | This paper | N/A |
| Human: HEK293T-TRIM21_N71D | This paper | N/A |
| Human: HEK293T-TRIM21_S80E | This paper | N/A |
| Human: HEK293T-TRIM21_M72E/S80E | This paper | N/A |
| Human: HEK293T-TRIM21_V43W | This paper | N/A |
| Human: HEK293T-TRIM21_V43W/M72E | This paper | N/A |
| Human: HEK293T-TRIM21_W112R | This paper | N/A |
| Human: HEK293T-TRIM21_V113E | This paper | N/A |
| Human: HEK293T-TRIM21_Q116R | This paper | N/A |
| Human: HEK293T-TRIM21_R118E | This paper | N/A |
| Human: HEK293T-TRIM21_E129A | This paper | N/A |
| Human: HEK293T-TRIM21_ΔBox | This paper | N/A |
| Human: HEK293T-mCherry-hTRIM21 | (Clift et al., 2017) | https://doi.org/10.1016/j.cell.2017.10.033 |
| Human: RPE-1 | ATCC | Cat#CRL-4000,: RRID:CVCL_4388 |
| Human: RPE-1 TRIM21 KO | This paper | N/A |
| Drosophila S2-DGRC | Drosophila Genomics Resource Centre | Cat#S2-DGRC |
| **Experimental Models: Organisms/Strains** |  |  |
| Mouse: CAG:H2B-EGFP | The Jackson Laboratory | Cat#JAX:006069; RRID:IMSR_JAX:006069 |
| **Recombinant DNA** |  |  |
| pGEMHE | (Liman et al., 1992a) | DOI: 10.1016/0896-6273(92)90239-a |
| pGEMHE-mEGFP | (Clift et al., 2017) | https://doi.org/10.1016/j.cell.2017.10.033 |
| pGEMHE-membrane-mEGFP | (Clift et al., 2017) | https://doi.org/10.1016/j.cell.2017.10.033 |
| pGEMHE-4xmyc-mEGFP (4myc) | This paper | N/A |
| pGEMHE-4xmyc-mEGFP (3myc) | This paper | N/A |
| pGEMHE-4xmyc-mEGFP (2myc) | This paper | N/A |
| pGEMHE-4xmyc-mEGFP (1myc) | This paper | N/A |
| pGEMHE-4xmyc-mEGFP (0myc) | This paper | N/A |
| pGEMHE-TRIM21-CRY2olig-mCh | This paper | N/A |
| pGEMHE-T21-M10E/M72E-CRY2olig-mCh | This paper | N/A |
| pGEMHE-TRIM21-E13A-CRY2olig-mCh | This paper | N/A |
| pGEMHE-TRIM21_RING-CRY2olig-mCh | This paper | N/A |
| pGEMHE-TRIM21_RB-CRY2olig-mCh | This paper | N/A |
| pGEMHE-TRIM21_RBCC-CRY2olig-mCh | This paper | N/A |
| pGEMHE-TRIM21_RB_S80A-CRY2olig-mCh | This paper | N/A |
| pGEMHE-TRIM21_RB_S80E-CRY2olig-mCh | This paper | N/A |
| pGEMHE-vhhGFP4-hlgG1-Fc | (Clift et al., 2017) | https://doi.org/10.1016/j.cell.2017.10.033 |
| pGEMHE-vhhGFP4-hlgG1-Fc(H433A) | This paper | N/A |
| pGEMHE-TRIM21 | This paper | N/A |
| pGEMHE-T21RBCC-vhhGFP4 | This paper | N/A |
| pGEMHE-T21RB-vhhGFP4 | This paper | N/A |
| pGEMHE-T21R-vhhGFP4 | This paper | N/A |
| pGEMHE-mCherry-T21RBCC-vhhGFP4 | This paper | N/A |
| pGEMHE- mCherry-T21RB-vhhGFP4 | This paper | N/A |
| pGEMHE- mCherry-T21R-vhhGFP4 | This paper | N/A |
| pT21MPP | (Kiss et al., 2019) | DOI: doi.org/10.1038/s41467-019-12388-y |
| pT21MPP -hT21_E12A | (Kiss et al., 2019) | DOI: doi.org/10.1038/s41467-019-12388-y |
| pT21MPP -hT21_E12R | (Kiss et al., 2019) | DOI: doi.org/10.1038/s41467-019-12388-y |
| pT21MPP -hT21_E12Y | (Kiss et al., 2019) | DOI: doi.org/10.1038/s41467-019-12388-y |
| pT21MPP -hT21_E13A | (Kiss et al., 2019) | DOI: doi.org/10.1038/s41467-019-12388-y |
| pT21MPP -hT21_E13R | (Kiss et al., 2019) | DOI: doi.org/10.1038/s41467-019-12388-y |
| pT21MPP -hT21_M10E | This paper | N/A |
| pT21MPP -hT21_M72E | This paper | N/A |
| pT21MPP -hT21_R67A | This paper | N/A |
| pT21MPP -hT21_N71D | This paper | N/A |
| pT21MPP -hT21_S80E | This paper | N/A |
| pT21MPP -hT21_M72E/S80E | This paper | N/A |
| pT21MPP -hT21_V43W | This paper | N/A |
| pT21MPP -hT21_V43W/M72E | This paper | N/A |
| pT21MPP -hT21_W112R | This paper | N/A |
| pT21MPP -hT21_V113E | This paper | N/A |
| pT21MPP -hT21_Q116R | This paper | N/A |
| pT21MPP -hT21_R118E | This paper | N/A |
| pT21MPP -hT21_E129A | This paper | N/A |
| pT21MPP -hT21_ΔBox | This paper | N/A |
| pPMEZ | This paper | N/A |
| pPMEZ-H2B-mEGFP-FKBP | This paper | N/A |
| pCRV-Gag-Pol | Stuart Neil | N/A |
| pMD2.G | Didier Trono | Addgene Plasmid #12259 |
| pGL4.32[*luc2P*/NF-κB-RE/Hygro] | Promega | Cat#E8491 |
| pMT107 (CMVp-Octa-6His-Ub) | (Treier et al., 1994) | DOI: 10.1016/s0092-8674(94)90502-9 |
| pcDNA3 | Invitrogen | Cat#V79020 |
| pcDNA3-2x-6His-Ub | This paper | N/A |
| pcDNA3-2x-6His-Ub_K48R | This paper | N/A |
| pcDNA3-2x-6His-Ub_K63R | This paper | N/A |
| pcDNA3-2x-6His-Ub_K48 only | This paper | N/A |
| pcDNA3-2x-6His-Ub_K63 only | This paper | N/A |
| pHR-CRY2Clust | This paper | N/A |
| pHR-CRY2Clust-TRIM21 | This paper | N/A |
| pH-vhhGFP4-RFP-CRY2Clust-TRIM21 | This paper | N/A |
| pHW-CIBN-MP_HRW-CRY2-V<sub>HH</sub> | (Osswald et al., 2019) | DOI: 10.3390/biom9020061 |
| pHGW-aPKC | (Osswald et al., 2019) | DOI: 10.3390/biom9020061 |
| MBP-T21CC (MBPCC235) | This paper | N/A |
| T21CC (CC235) | This paper | N/A |
| pOPTG | (Ghazi-Tabatabai et al., 2008) | DOI: 10.1016/j.str.2008.06.010 |
| pOPTG- TRIM21_RL | (Dickson et al., 2018) | DOI: 10.7554/eLife.32660 |
| pOPTG- TRIM21_RB | Dickson et al., 2018) | DOI: 10.7554/eLife.32660 |
| pOPTG- TRIM21_RLR | This study | N/A |
| pOPTG- TRIM21_RLR_M10E/M10E | This study | N/A |
| pOPTG- TRIM21_RLR_N71D/N71D | This study | N/A |
| pOPTG- TRIM21_RBCC | This study | N/A |
| pFastBac-MBP-TRIM21 | This study | N/A |
| pOPTH | (Ghazi-Tabatabai et al., 2008) | DOI: 10.1016/j.str.2008.06.010 |
| pOPTH-T21R-vhhGFP4 | This study | N/A |
| pOPTH-T21RB-vhhGFP4 | This study | N/A |
| **Software and Algorithms** |  |  |
| Prism | GraphPad Software | N/A |
| FlowJo | FLOWJO, LLC | N/A |
| ImageJ/FIJI | (Schindelin et al., 2012) | doi.org/10.1038/nmeth.2019 |
| IncuCyte software | Sartorius | N/A |
| **Other** |  |  |
| Leica SP8 microscope | Leica | N/A |
| Nikon Ti microscope | Nikon | N/A |
| Dark Reader Transilluminator | Clare Chemical Research | Cat#DR46B |

## REFERENCES

Baek, K., Krist, D.T., Prabu, J.R., Hill, S., Klügel, M., Neumaier, L.-M., von Gronau, S., Kleiger, G., and Schulman, B.A. (2020). NEDD8 nucleates a multivalent cullin–RING– UBE2D ubiquitin ligation assembly. Nature 578, 461–466.

Baudisch, B., Pfort, I., Sorge, E., and Conrad, U. (2018). Nanobody-Directed Specific Degradation of Proteins by the 26S-Proteasome in Plants. Frontiers in Plant Science 9.

Buetow, L., and Huang, D.T. (2016). Structural insights into the catalysis and regulation of E3 ubiquitin ligases. Nature Reviews Molecular Cell Biology 17, 626–642.

Bugaj, L.J., Choksi, A.T., Mesuda, C.K., Kane, R.S., and Schaffer, D.V. (2013). Optogenetic protein clustering and signaling activation in mammalian cells. Nature Methods 10, 249–252.

Campbell, R.E., Tour, O., Palmer, A.E., Steinbach, P.A., Baird, G.S., Zacharias, D.A., and Tsien, R.Y. (2002). A monomeric red fluorescent protein. Proceedings of the National Academy of Sciences 99, 7877.

Castro-Dopico, T., Dennison, T.W., Ferdinand, J.R., Mathews, R.J., Fleming, A., Clift, D., Stewart, B.J., Jing, C., Strongili, K., Labzin, L.I., et al. (2019). Anti-commensal IgG Drives Intestinal Inflammation and Type 17 Immunity in Ulcerative Colitis. Immunity 50, 1099–1114 e1010.

Caussinus, E., Kanca, O., and Affolter, M. (2012). Fluorescent fusion protein knockout mediated by anti-GFP nanobody. Nature Structural & Molecular Biology 19, 117–121.

Chen, X., Liu, M., Lou, H., Lu, Y., Zhou, M.-T., Ou, R., Xu, Y., and Tang, K.-F. (2019). Degradation of endogenous proteins and generation of a null-like phenotype in zebrafish using Trim-Away technology. Genome Biology 20, 19.

Clift, D., McEwan, W.A., Labzin, L.I., Konieczny, V., Mogessie, B., James, L.C., and Schuh, M. (2017). A Method for the Acute and Rapid Degradation of Endogenous Proteins. Cell 171, 1692–1706.e1618.

Clift, D., So, C., McEwan, W.A., James, L.C., and Schuh, M. (2018). Acute and rapid degradation of endogenous proteins by Trim-Away. Nature Protocols 13, 2149–2175.

Cole, D., Young, G., Weigel, A., Sebesta, A., and Kukura, P. (2017). Label-Free Single-Molecule Imaging with Numerical-Aperture-Shaped Interferometric Scattering Microscopy. ACS Photonics 4, 211–216.

Coyne, A.N., Zaepfel, B.L., Hayes, L., Fitchman, B., Salzberg, Y., Luo, E.C., Bowen, K., Trost, H., Aigner, S., Rigo, F., et al. (2020). G4C2 Repeat RNA Initiates a POM121-Mediated Reduction in Specific Nucleoporins in C9orf72 ALS/FTD. Neuron.

Deng, W., Bates, J.A., Wei, H., Bartoschek, M.D., Conradt, B., and Leonhardt, H. (2020). Tunable light and drug induced depletion of target proteins. Nature Communications 11, 304.

Deshaies, R.J., and Joazeiro, C.A.P. (2009). RING Domain E3 Ubiquitin Ligases. Annual Review of Biochemistry 78, 399–434.

Dickson, C., Fletcher, A.J., Vaysburd, M., Yang, J.-C., Mallery, D.L., Zeng, J., Johnson, C.M., McLaughlin, S.H., Skehel, M., Maslen, S., et al. (2018). Intracellular antibody signalling is regulated by phosphorylation of the Fc receptor TRIM21. eLife 7, e32660.

Esposito, D., Koliopoulos, M.G., and Rittinger, K. (2017). Structural determinants of TRIM protein function. Biochemical Society Transactions 45, 183–191.

Fan, W., Zhang, D., Qian, P., Qian, S., Wu, M., Chen, H., and Li, X. (2016). Swine TRIM21 restricts FMDV infection via an intracellular neutralization mechanism. Antiviral Res 127, 32–40.

Fant, C.B., Levandowski, C.B., Gupta, K., Maas, Z.L., Moir, J., Rubin, J.D., Sawyer, A., Esbin, M.N., Rimel, J.K., Luyties, O., et al. (2020). TFIID Enables RNA Polymerase II Promoter-Proximal Pausing. Mol Cell 78, 785–793 e788.

Fletcher, A.J., Mallery, D.L., Watkinson, R.E., Dickson, C.F., and James, L.C. (2015). Sequential ubiquitination and deubiquitination enzymes synchronize the dual sensor and effector functions of TRIM21. Proc Natl Acad Sci U S A 112, 10014–10019.

Fletcher, A.J., Vaysburd, M., Maslen, S., Zeng, J., Skehel, J.M., Towers, G.J., and James, L.C. (2018). Trivalent RING Assembly on Retroviral Capsids Activates TRIM5 Ubiquitination and Innate Immune Signaling. Cell Host Microbe 24, 761–775 e766.

Foss, S., Bottermann, M., Jonsson, A., Sandlie, I., James, L.C., and Andersen, J.T. (2019). TRIM21—From Intracellular Immunity to Therapy. Frontiers in Immunology 10.

Foss, S., Watkinson, R.E., Grevys, A., McAdam, M.B., Bern, M., Høydahl, L.S., Dalhus, B., Michaelsen, T.E., Sandlie, I., James, L.C., et al. (2016). TRIM21 Immune Signaling Is More Sensitive to Antibody Affinity Than Its Neutralization Activity. The Journal of Immunology 196, 3452.

Franke, D., Kikhney, A. G., & Svergun, D. I. (2012). Automated acquisition and analysis of small angle X-ray scattering data. Nuc Inst Meth A 689, 52–59.

Franke, D., and Svergun, D.I. (2009). DAMMIF, a program for rapid ab-initio shape determination in small-angle scattering. J Appl Crystallogr 42, 342–346.

Fulcher, L.J., Macartney, T., Bozatzi, P., Hornberger, A., Rojas-Fernandez, A., and Sapkota, G.P. (2016). An affinity-directed protein missile system for targeted proteolysis. Open Biol 6.

Ghazi-Tabatabai, S., Saksena, S., Short, J.M., Pobbati, A.V., Veprintsev, D.B., Crowther, R.A., Emr, S.D., Egelman, E.H., and Williams, R.L. (2008). Structure and disassembly of filaments formed by the ESCRT-III subunit Vps24. Structure 16, 1345–1356.

Gross, G.G., Straub, C., Perez-Sanchez, J., Dempsey, W.P., Junge, J.A., Roberts, R.W., Trinh, L.A., Fraser, S.E., De Koninck, Y., De Koninck, P., et al. (2016). An E3-ligase-based method for ablating inhibitory synapses. Nature Methods 13, 673–678.

Hayer, A., Stoeber, M., Bissig, C., and Helenius, A. (2010). Biogenesis of Caveolae: Stepwise Assembly of Large Caveolin and Cavin Complexes. Traffic 11, 361–382.

Hilpert, K., Hansen, G., Wessner, H., Küttner, G., Welfle, K., Seifert, M., and Höhne, W. (2001). Anti-c-myc antibody 9E10: epitope key positions and variability characterized using peptide spot synthesis on cellulose. Protein Engineering, Design and Selection 14, 803–806.

Hofmann, R.M., and Pickart, C.M. (2001). In vitro assembly and recognition of Lys-63 polyubiquitin chains. J Biol Chem 276, 27936–27943.

Ibrahim, A.F.M., Shen, L., Tatham, M.H., Dickerson, D., Prescott, A.R., Abidi, N., Xirodimas, D.P., and Hay, R.T. (2020). Antibody RING-Mediated Destruction of Endogenous Proteins. Mol Cell.

Israel, S., Casser, E., Drexler, H.C.A., Fuellen, G., and Boiani, M. (2019). A framework for TRIM21-mediated protein depletion in early mouse embryos: recapitulation of Tead4 null phenotype over three days. BMC Genomics 20, 755.

Jacques, D.A., and Trewhella, J. (2010). Small-angle scattering for structural biology--expanding the frontier while avoiding the pitfalls. Protein Science 19, 642–657.

James, L.C., Keeble, A.H., Khan, Z., Rhodes, D.A., and Trowsdale, J. (2007). Structural basis for PRYSPRY-mediated tripartite motif (TRIM) protein function. Proc Natl Acad Sci U S A 104, 6200–6205.

Ju Shin, Y., Kyun Park, S., Jung Jung, Y., Na Kim, Y., Sung Kim, K., Kyu Park, O., Kwon, S.-H., Ho Jeon, S., Trinh, L.A., Fraser, S.E., et al. (2015). Nanobody-targeted E3-ubiquitin ligase complex degrades nuclear proteins. Scientific Reports 5, 14269.

Kanda, T., Sullivan, K.F., and Wahl, G.M. (1998). Histone-GFP fusion protein enables sensitive analysis of chromosome dynamics in living mammalian cells. Curr Biol 8, 377–385.

Kiss, L., Zeng, J., Dickson, C.F., Mallery, D.L., Yang, J.C., McLaughlin, S.H., Boland, A., Neuhaus, D., and James, L.C. (2019). A tri-ionic anchor mechanism drives Ube2N-specific recruitment and K63-chain ubiquitination in TRIM ligases. Nat Commun 10, 4502.

Konarev, P.V., Volkov, V.V., Sokolova, A.V., Koch, M.H.J., and Svergun, D.I. (2003). PRIMUS - a Windows-PC based system for small-angle scattering data analysis. J Appl Cryst 36, 1277–1282.

Krauss, N., Wessner, H., Welfle, K., Welfle, H., Scholz, C., Seifert, M., Zubow, K., Ay, J., Hahn, M., Scheerer, P., et al. (2008). The structure of the anti-c-myc antibody 9E10 Fab fragment/epitope peptide complex reveals a novel binding mode dominated by the heavy chain hypervariable loops. Proteins 73, 552–565.

Lim, S., Khoo, R., Peh, K.M., Teo, J., Chang, S.C., Ng, S., Beilhartz, G.L., Melnyk, R.A., Johannes, C.W., Brown, C.J., et al. (2020). bioPROTACs as versatile modulators of intracellular therapeutic targets including proliferating cell nuclear antigen (PCNA). Proceedings of the National Academy of Sciences 117, 5791.

Liman, E.R., Tytgat, J., and Hess, P. (1992a). Subunit stoichiometry of a mammalian K+ channel determined by construction of multimeric cDNAs. Neuron 9, 861–871.

Liman, E.R., Tytgat, J., and Hess, P. (1992b). Subunit stoichiometry of a mammalian K+ channel determined by construction of multimeric cDNAs. Neuron 9, 861–871.

Ludwicki, M.B., Li, J., Stephens, E.A., Roberts, R.W., Koide, S., Hammond, P.T., and DeLisa, M.P. (2019). Broad-Spectrum Proteome Editing with an Engineered Bacterial Ubiquitin Ligase Mimic. ACS Central Science 5, 852–866.

Mallery, D.L., McEwan, W.A., Bidgood, S.R., Towers, G.J., Johnson, C.M., and James, L.C. (2010). Antibodies mediate intracellular immunity through tripartite motif-containing 21 (TRIM21). Proceedings of the National Academy of Sciences 107, 19985–19990.

Marschall, A.L.J., Single, F.N., Schlarmann, K., Bosio, A., Strebe, N., van den Heuvel, J., Frenzel, A., and Dübel, S. (2014). Functional knock down of VCAM1 in mice mediated by endoplasmatic reticulum retained intrabodies. mAbs 6, 1394–1401.

Más, P., Devlin, P.F., Panda, S., and Kay, S.A. (2000). Functional interaction of phytochrome B and cryptochrome 2. Nature 408, 207–211.

McEwan, W.A., Falcon, B., Vaysburd, M., Clift, D., Oblak, A.L., Ghetti, B., Goedert, M., and James, L.C. (2017). Cytosolic Fc receptor TRIM21 inhibits seeded tau aggregation. Proceedings of the National Academy of Sciences, 201607215.

McEwan, W.A., Hauler, F., Williams, C.R., Bidgood, S.R., Mallery, D.L., Crowther, R.A., and James, L.C. (2012). Regulation of Virus Neutralization and the Persistent Fraction by TRIM21. Journal of Virology 86, 8482–8491.

McEwan, W.A., Mallery, D.L., Rhodes, D.A., Trowsdale, J., and James, L.C. (2011). Intracellular antibody-mediated immunity and the role of TRIM21. Bioessays 33, 803–809.

McEwan, W.A., Tam, J.C.H., Watkinson, R.E., Bidgood, S.R., Mallery, D.L., and James, L.C. (2013). Intracellular antibody-bound pathogens stimulate immune signaling via the Fc receptor TRIM21. Nature Immunology 14, 327–336.

Mund, T., and Pelham, H.R. (2018). Substrate clustering potently regulates the activity of WW-HECT domain-containing ubiquitin ligases. J Biol Chem 293, 5200–5209.

Napolitano, L.M., and Meroni, G. (2012). TRIM family: Pleiotropy and diversification through homomultimer and heteromultimer formation. IUBMB Life 64, 64–71.

Nicolau, C.A., Gavard, J., and Bidere, N. (2020). TAK1 lessens the activity of the paracaspase MALT1 during T cell receptor signaling. Cell Immunol 353, 104115.

Ohtake, F., Tsuchiya, H., Saeki, Y., and Tanaka, K. (2018). K63 ubiquitylation triggers proteasomal degradation by seeding branched ubiquitin chains. Proceedings of the National Academy of Sciences 115, E1401.

Osswald, M., Santos, A.F., and Morais-de-Sa, E. (2019). Light-Induced Protein Clustering for Optogenetic Interference and Protein Interaction Analysis in Drosophila S2 Cells. Biomolecules 9.

Park, H., Kim, N.Y., Lee, S., Kim, N., Kim, J., and Heo, W.D. (2017). Optogenetic protein clustering through fluorescent protein tagging and extension of CRY2. Nature Communications 8, 30.

Portnoff, A.D., Stephens, E.A., Varner, J.D., and DeLisa, M.P. (2014). Ubiquibodies, synthetic E3 ubiquitin ligases endowed with unnatural substrate specificity for targeted protein silencing. J Biol Chem 289, 7844–7855.

Rakebrandt, N., Lentes, S., Neumann, H., James, L.C., and Neumann-Staubitz, P. (2014). Antibody- and TRIM21-dependent intracellular restriction of Salmonella enterica. Pathogens and Disease 72, 131–137.

Ravetch, J.V., and Bolland, S. (2001). IgG Fc Receptors. Annual Review of Immunology 19, 275–290.

Saeki, Y., Kudo, T., Sone, T., Kikuchi, Y., Yokosawa, H., Toh-e, A., and Tanaka, K. (2009). Lysine 63-linked polyubiquitin chain may serve as a targeting signal for the 26S proteasome. EMBO J 28, 359–371.

Saerens, D., Pellis, M., Loris, R., Pardon, E., Dumoulin, M., Matagne, A., Wyns, L., Muyldermans, S., and Conrath, K. (2005). Identification of a universal VHH framework to graft non-canonical antigen-binding loops of camel single-domain antibodies. J Mol Biol 352, 597–607.

Schapira, M., Calabrese, M.F., Bullock, A.N., and Crews, C.M. (2019). Targeted protein degradation: expanding the toolbox. Nat Rev Drug Discov 18, 949–963.

Schindelin, J., Arganda-Carreras, I., Frise, E., Kaynig, V., Longair, M., Pietzsch, T., Preibisch, S., Rueden, C., Saalfeld, S., Schmid, B., et al. (2012). Fiji: an open-source platform for biological-image analysis. Nat Methods 9, 676–682.

Schrader, E.K., Harstad, K.G., and Matouschek, A. (2009). Targeting proteins for degradation. Nature Chemical Biology 5, 815–822.

Shvets, E., Bitsikas, V., Howard, G., Hansen, C.G., and Nichols, B.J. (2015). Dynamic caveolae exclude bulk membrane proteins and are required for sorting of excess glycosphingolipids. Nature Communications 6, 6867.

So, C., Seres, K.B., Steyer, A.M., Mönnich, E., Clift, D., Pejkovska, A., Möbius, W., and Schuh, M. (2019). A liquid-like spindle domain promotes acentrosomal spindle assembly in mammalian oocytes. Science 364, eaat9557.

Stewart, M.D., Duncan, E.D., Coronado, E., DaRosa, P.A., Pruneda, J.N., Brzovic, P.S., and Klevit, R.E. (2017). Tuning BRCA1 and BARD1 activity to investigate RING ubiquitin ligase mechanisms. Protein Sci 26, 475–483.

Svergun, D.I. (1992). Determination of the regularization parameter in indirect-transform methods using perceptual criteria. J Appl Crystallogr 25, 495–503.

Svergun, D.I., Barberato, C., and Koch, M.H.J. (1995). CRYSOL - a Program to Evaluate X-ray Solution Scattering of Biological Macromolecules from Atomic Coordinates. J Appl Cryst 28, 768–773.

Taslimi, A., Vrana, J.D., Chen, D., Borinskaya, S., Mayer, B.J., Kennedy, M.J., and Tucker, C.L. (2014a). An optimized optogenetic clustering tool for probing protein interaction and function. Nature Communications 5, 4925.

Taslimi, A., Vrana, J.D., Chen, D., Borinskaya, S., Mayer, B.J., Kennedy, M.J., and Tucker, C.L. (2014b). An optimized optogenetic clustering tool for probing protein interaction and function. Nature Communications 5, 4925.

Traub, L.M. (2019). A nanobody-based molecular toolkit provides new mechanistic insight into clathrin-coat initiation. eLife 8, e41768.

Treier, M., Staszewski, L.M., and Bohmann, D. (1994). Ubiquitin-dependent c-Jun degradation in vivo is mediated by the delta domain. Cell 78, 787–798.

Vaysburd, M., Watkinson, R.E., Cooper, H., Reed, M., O’Connell, K., Smith, J., Cruickshanks, J., and James, L.C. (2013). Intracellular antibody receptor TRIM21 prevents fatal viral infection. Proceedings of the National Academy of Sciences 110, 12397.

Verma, R., Mohl, D., and Deshaies, R.J. (2020). Harnessing the Power of Proteolysis for Targeted Protein Inactivation. Mol Cell 77, 446–460.

Wagner, J.M., Roganowicz, M.D., Skorupka, K., Alam, S.L., Christensen, D., Doss, G., Wan, Y., Frank, G.A., Ganser-Pornillos, B.K., Sundquist, W.I., et al. (2016). Mechanism of B-box 2 domain-mediated higher-order assembly of the retroviral restriction factor TRIM5α. eLife 5, e16309.

Watkinson, R.E., McEwan, W.A., Tam, J.C., Vaysburd, M., and James, L.C. (2015). TRIM21 Promotes cGAS and RIG-I Sensing of Viral Genomes during Infection by Antibody-Opsonized Virus. PLoS Pathog 11, e1005253.

Wu, T., Yoon, H., Xiong, Y., Dixon-Clarke, S.E., Nowak, R.P., and Fischer, E.S. (2020). Targeted protein degradation as a powerful research tool in basic biology and drug target discovery. Nature Structural & Molecular Biology.

Yamada, T., Yang, Y., and Bonni, A. (2013). Spatial organization of ubiquitin ligase pathways orchestrates neuronal connectivity. Trends Neurosci 36, 218–226.

Yamaguchi, N., Colak-Champollion, T., and Knaut, H. (2019). zGrad is a nanobody-based degron system that inactivates proteins in zebrafish. eLife 8, e43125.

Young, G., Hundt, N., Cole, D., Fineberg, A., Andrecka, J., Tyler, A., Olerinyova, A., Ansari, A., Marklund, E.G., Collier, M.P., et al. (2018). Quantitative mass imaging of single biological macromolecules. Science 360, 423–427.

Zeng, J., Slodkowicz, G., and James, L.C. (2019). Rare missense variants in the human cytosolic antibody receptor preserve antiviral function. eLife 8, e48339.

Zheng, N., and Shabek, N. (2017). Ubiquitin Ligases: Structure, Function, and Regulation. Annu Rev Biochem 86, 129–157.

